# A neural substrate for resistance to change in the ventral hippocampus

**DOI:** 10.64898/2026.04.22.720018

**Authors:** Aika Saito, Hiroyoshi Ogishima, Asuka Joji-Nishino, Kazuo Emoto, Saori C Tanaka, Akira Uematsu

**Author notes:** correspondence should be addressed to A.U.

## Abstract

Strong prior beliefs can render individuals resistant to change, even when outcomes contradict expectations, yet the neural mechanisms supporting such stability remain poorly understood. Here we show that the ventral hippocampus (vCA1) actively stabilizes inferred contextual state (belief) during aversive memory extinction in a behavior-dependent manner. Using *in vivo* calcium imaging, we identify a subset of vCA1 neurons that exhibit selective reductions in activity when expected aversive outcomes do not occur during extinction. Axonal calcium imaging further reveals that reduced signals are broadcast to basal amygdala, nucleus accumbens, and medial prefrontal cortex. Optogenetic manipulations demonstrate that these signals actively suppress extinction learning and promote persistence of defensive response. Computational modeling indicates that vCA1 omission-related signals reflect discrepancies between beliefs and observed omission of the expected outcome, thereby maintaining the aversive belief. Crucially, convergent evidence from calcium imaging, modelling, and closed-loop optogenetic manipulations reveals that vCA1 omission-related signals are specific to the period during which animals execute defensive behavior driven by their inferred aversive state. These findings identify a novel mechanism in vCA1 that stabilizes inferred aversive state despite changes in stimulus contingencies and suggest a key role of vCA1 in belief-based control of learning.

## INTRODUCTION

Although prediction error theories posit that behavior is updated in response to new evidence ^1-7^, in real behavior this is often not the case. Individuals also rely on memories of particularly salient experiences to preserve specific behavioral patterns and their associated predictions, maintaining an inferred contextual state (i.e., belief) even in the face of new information. Indeed, strong prior beliefs have been shown to promote resistance to contradictory information ^8,9^. Clinical observations further suggest that symptoms associated with mental disorders are linked to excessive maintenance of prior expectation, as seen in phenomena such as cognitive immunization and negative expectancy bias ^10,11^. Despite these observations, the neural mechanisms that mediate maintenance of prior belief and behavior remain poorly understood.

The hippocampus is proposed as a neural substrate for inferring latent causes, or latent contextual states, that explain sensory experiences ^12^. According to the latent cause model, the hippocampus clusters experiences into distinct contexts, such that new learning reflects assigning new experiences to a separate latent cause (e.g., “I may be in danger” vs. “I may be in safe”). A recent study supports the key role of hippocampus in the latent state inference during decision-making process ^13^. Beyond these findings, the hippocampus switches between memory encoding and retrieval by computing the overlap between expected and encountered stimuli ^14-17^, suggesting that it continually compares internal predictions with sensory input. Furthermore, it has been shown that pre-existing hippocampal network dynamics constrain the formation of new representations ^18^. Thus, the hippocampus may compute discrepancies between inferred state based on previously encoded memory and new experiences, biasing the system toward maintaining previously encoded state rather than driving new learning.

## RESULTS

### vCA1 responds to the omission of an expected aversive outcome

Given that the ventral CA1 (vCA1) region encode nonspatial episodic information such as past reward and aversive experiences ^19-21^, we hypothesized that vCA1 would represent error signals linked to prior experiences when a non-spatial stimulus contingency changes. To investigate our hypothesis, we performed in vivo Ca^2+^ imaging of vCA1 neurons in freely moving mice using a miniature microscope. Mice first received an injection of adeno-associated virus (AAV) expressing jGCaMP8s followed by GRIN lens implantation (**Figure 1a**). Because aversive memories are known to resist modification ^22,23^, we employed aversive memory paradigms. Mice were subjected to differential aversive conditioning, in which one initially neutral tone (conditioned stimulus; CS^+^) was paired with a footshock (unconditioned stimulus; US) while a second tone (CS^-^) was not (**Figure 1b**). On the next day, mice were exposed to repeated CS^+^ exposure without foot shock during an extinction session in which errors are generated due to the omission of the expected shock. We confirmed that mice acquired aversive and extinction memory after these sessions (**Figure 1c**).

**Figure 1.**
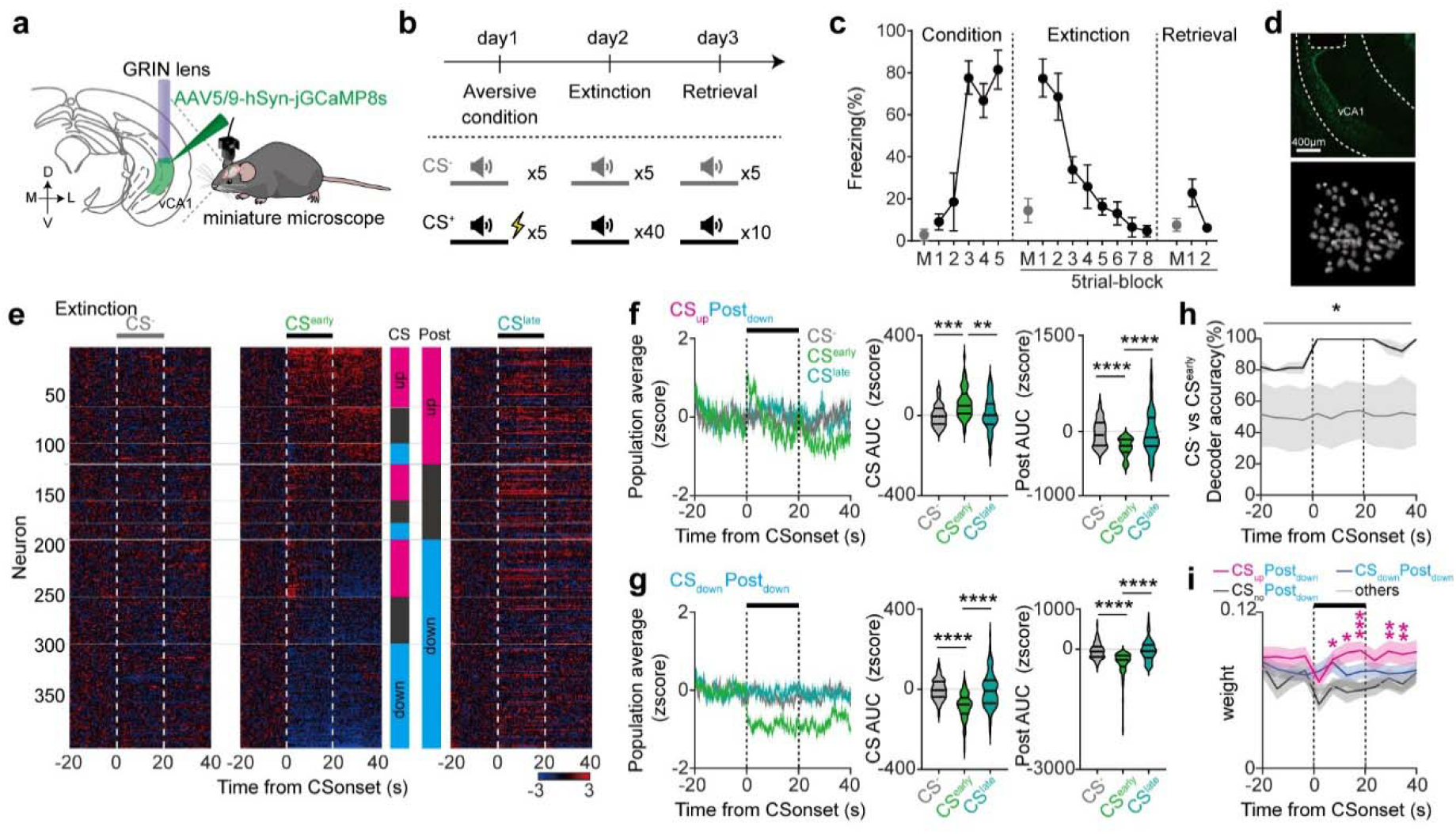
vCA1 neurons signal omission of expected outcomes during aversive extinction. **(a)** In-vivo single cell imaging from vCA1. AAV encoding jGCaMP8s was injected into vCA1 and a GRIN lens was implanted above the injection site. **(b)** Aversive conditioning and extinction paradigm. **(c)** Percentage of freezing exhibited during CS^-^ (M) and CS^+^ (n = 5). **(d)** jGCaMP8s expression and GRIN lens implantation in vCA1. Scale bar, 400μm (top). Maximum projection image of vCA1 neurons acquired with a miniscope (bottom). **(e)** Heat maps of calcium activity triggered by CS onset during CS^-^(left), the CS^early^ (middle) and CS^late^ (right) phases in extinction session (n = 398 cells, 5 mice). Magenta and blue squares indicate up and down neurons, respectively. CS_up_Post_up_ (n = 59), CS_no_Post_up_ (n = 36), CS_down_Post_up_ (n = 21), CS_up_Post_no_ (n = 36), CS_no_Post_no_ (n = 23), CS_down_Post_no_ (n = 16), CS_up_ Post_down_ (n = 57), CS_no_Post_down_ (n = 46), and CS_down_Post_down_ (n = 104) groups. **(f)** Population activity of CS_up_Post_down_ cells during CS^-^, CS^early^ and CS^late^ of extinction (left). Mean AUC of CS_up_Post_down_ cells revealed a significantly higher response during CS^early^ compared to CS^−^ (middle), and a decreased response during the post-CS period of CS^early^ compared to CS^-^ (right). **(g)** Population activity of CS_down_Post_down_ cells during CS^-^, CS^early^ and CS^late^ of extinction (left). Mean AUC of CS_down_Post_down_ cells revealed a significantly lower response during CS^early^ (middle) and post-CS (right) compared to CS^−^. **(h)** Time-resolved decoding accuracy of CS^-^ and CS^early^ identity. Decoding accuracy exceeded that of label-shuffled data (gray line) at all time points. **(i)** Weights of CS_up_Post_down_, CS_no_Post_down_, CS_down_Post_down,_ and others in linear decoder as a function of time. CS_up_Post_down_ cells showed the highest classifier weights during CS and post-CS periods. \**P* < 0.05; \*\**P* < 0.01; \*\*\**P* < 0.001; \*\*\*\**P* < 0.0001; For statistical details, see Supplementary Table 1.

We collected calcium imaging data from a total of 398 neurons across 5 mice during the extinction session (**Figure 1d; Extended Data Fig. 1a**). We analyzed neural responses at the CS and post-CS period during the early phase of extinction (CS^early^) to assess cue-related responses and responses to expected shock omission, respectively. Based on the direction of responses (up, down, or no change) in each period, vCA1 neurons were classified into nine functional clusters representing all possible combinations of CS and post-CS responses (**Figure 1e-g; Extended Data Fig. 1b-f**). Among all recorded neurons, 15% exhibited excitatory CS-evoked responses followed by sustained excitatory post-CS activity (CS_up_Post_up_). A smaller fraction (5%) showed inhibitory CS-evoked responses followed by excitatory post-CS responses (CS_down_Post_up_). Fourteen percent of neurons displayed excitatory CS responses followed by inhibitory post-CS responses (CS_up_Post_down_), and the largest fraction (26%) exhibited inhibitory responses during both the CS and post-CS periods (CS_down_Post_down_). At the population level, CS and post-CS responses of these 4 populations were significantly higher or lower in CS^early^ than in CS^-^, reflecting aversive learning induced CS response and response associated with the omission of expected shock, respectively. In addition, CS_up_Post_down_ and CS_down_Post_up_ displayed opposite polarities to the CS and shock omission during early period of extinction similar to prediction error signals seen in dopamine neurons ^24,25^. However, unlike the rapid onset of predicted shock omission response in dopamine neurons ^24,25^, those in CS_up_Post_down_ and CS_down_Post_up_ neurons exhibited slow transient (see Supplementary Table 2), suggesting that these error signals are distinct from aversive dopaminergic prediction error.

Taking advantage of across-day cell tracking enabled by the miniature microscope, we next quantified shock-induced responses which had occurred during prior aversive conditioning (**Extended Data Fig. 2a-c**). Across most functional clusters, more than 40% of neurons exhibited significant shock-evoked responses, except for CS_down_Post_up_ neurons (**Extended Data Fig. 2d**). Comparison across conditioning trials revealed little to no change in the magnitude of these shock responses (**Extended Data Fig. 2e-h**), indicating that shock-evoked activity was largely stable and did not undergo substantial trial-by-trial modulation. We then investigated whether each subtype develops CS responses during the association with aversive outcomes. The CS_up_Post_down_ subtype exhibited stronger CS response in the last trial of conditioning (**Extended Data Fig. 2e**), indicating that CS_up_Post_down_ subtype developed CS-evoked response during CS-US association and that CS responses persisted into the early phase of extinction. Together, these data show that while cells respond to different task related events, specific subsets of vCA1 neurons not only respond to aversive shock but also acquire CS-evoked responses during conditioning. During extinction, these neurons exhibit CS-evoked activity and prolonged reduction upon the omission of expected shock.

### CS_up_Post_down_ neurons drive population level omission coding

To investigate if vCA1 neuronal population represents aversive-predictive cues and shock omissions in a manner distinct from neutral cues and non-predictive trials, we applied a population decoding approach. Mean calcium activity from the recorded neurons across all mice at a given time point was compared for CS^-^ and CS^early^ (**Extended Data Fig. 1h**). We obtained high decoding accuracy across all time points with CS and post-CS periods having higher accuracy than baseline (**Figure 1h**). To evaluate the contribution of functional neuronal groups to the linear decoder, we examined the decoder weights at each time point. We observed that CS_up_Post_down_ cells had the highest weights during the CS and post-CS periods (**Figure 1i; Extended Data Fig. 1i**), suggesting that CS_up_Post_down_ cells play a critical role in encoding CS and expected shock omission signals at the population level.

### vCA1 neurons exhibit reduced responses to the omission of an expected appetitive outcome

The observed post-CS suppression following shock omission may reflect the positive valence of unexpected safety. To test whether this suppression is driven by outcome valence or by the omission of expected outcome independent of valence, we examined neuronal responses during omission of an expected reward. If suppression reflects value-based coding, omission of an appetitive outcome should not elicit a similar response pattern. Following the aversive conditioning paradigms, several mice were trained on appetitive conditioning followed by an omission session. A distinct auditory tone (CS^r^) was paired with sucrose delivery for 2 consecutive days. On the next day, mice underwent an omission paradigm in which sucrose delivery was omitted on 20% of all trials in a randomized manner (**Extended Data Fig. 3a**). Mice displayed robust anticipatory licking during the CS period in both sucrose and omission trials, but licking was rapidly decreased following CS offset in omission trials (**Extended Data Fig. 3a**).

We collected calcium signals from a total of 131 neurons across 3 mice. At the population level, CS responses were comparable between reward and omission trials, whereas post-CS activity was significantly lower during sucrose omission compared with sucrose delivery. Notably, 13% of recorded neurons exhibited increased activity during the CS period and suppressed activity after sucrose omission period. These CS_up_Post_down_ neurons displayed increased post-CS responses when sucrose was delivered (**Extended Data Fig. 3b-e**).

Similar to aversive extinction, we next tested whether vCA1 neuronal population could discriminate appetitive-predictive cues and sucrose presentation from same cue and sucrose omission using a population decoding approach (**Extended Data Fig. 3f**). Decoding accuracy significantly increased during the early and late post-CS periods but remained at chance level during the CS period (**Extended Data Fig. 3g**). CS_up_Post_down_ neurons carried higher decoder weights than other functional types, similar to our observations during aversive extinction (**Extended Data Fig. 3h**). To further examine the contribution of highly weighted neurons in linear decoder, we analyzed the activity of the top 20% of weights. These neurons exhibited the reduced post-CS activity in omission trials relative to sucrose trials (**Extended Data Fig. 3i**), suggesting that omission-related suppression is a major contributor to population decoding. Overall, vCA1 neurons show reduced activity in response to the expected outcome omission independent of valence, which contributes to population level decoding.

### vCA1 broadcasts omission-related reduced activity to multiple brain regions

Because vCA1 harbors highly segregated neuronal populations defined by projection targets and distinct vCA1 projections regulate different aspects of emotional memory ^20,26-28^, we next sought to identify how vCA1 sends omission response to multiple output regions during aversive extinction. Among the various target regions of the vCA1, we focused on the ventromedial prefrontal cortex (vmPFC), the nucleus accumbens (NAc), and the basal amygdala (BA), which have been demonstrated to play a pivotal role in aversive memory extinction ^29-34^. We injected an AAV encoding a Ca^2+^ indicator (jRGECO1a) into the vCA1 and implanted optic fibers into the vmPFC, NAc, and BA (**Figure 2a; Extended Data Fig. 4a**). Using a multi-fiber photometry system, we simultaneously monitored the bulk terminal activities of the vmPFC, NAc, and BA (**Figure 2a**). Similar to the miniature microscope study, mice were subjected to a series of aversive conditioning, extinction, and retrieval experiments. At the behavior level, mice acquired both aversive memory and extinction successfully (**Figure 2b, c**).

**Figure 2.**
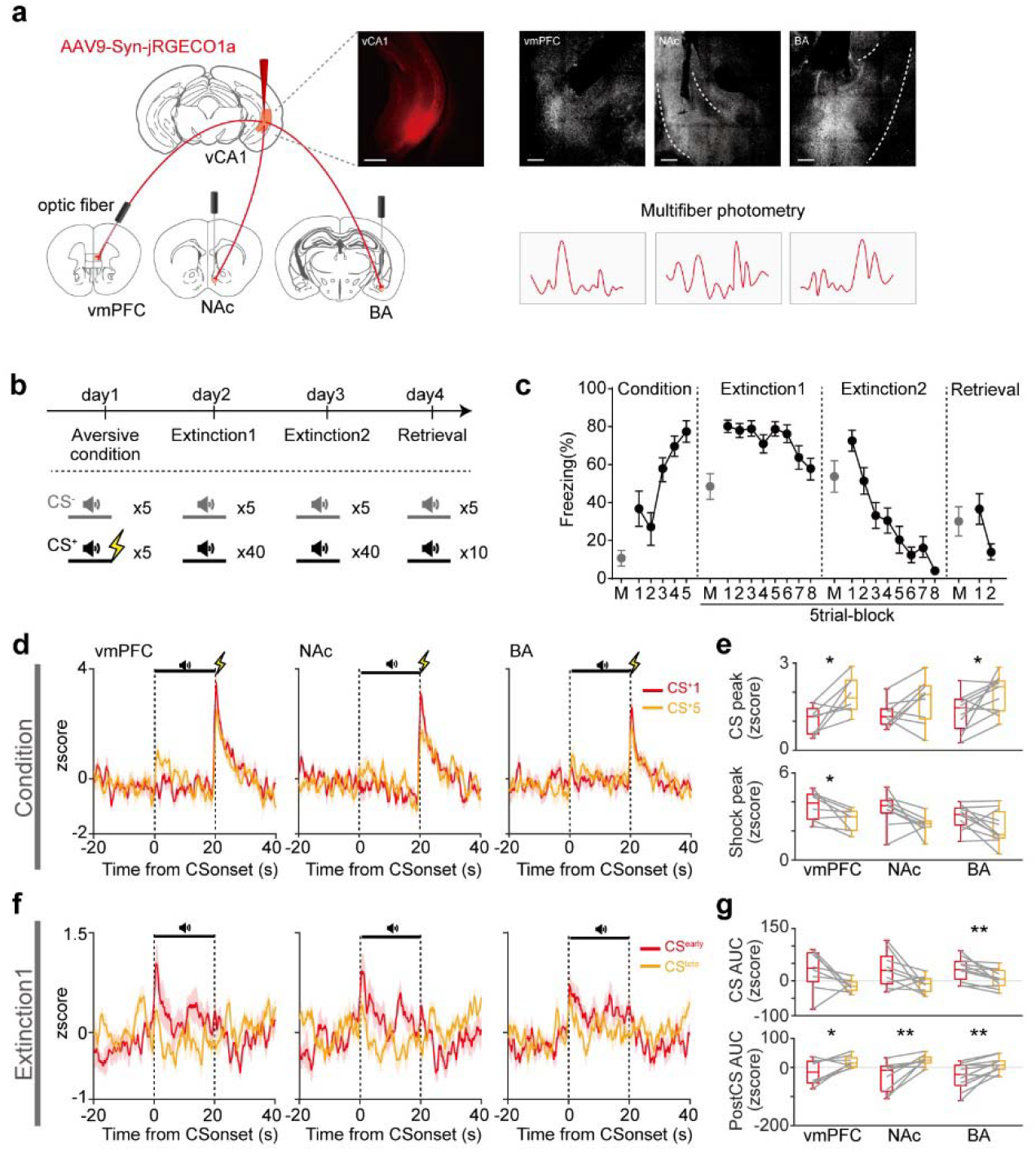
Calcium activity in vCA1 axon terminals exhibits omission-induced suppression. **(a)** Simultaneous monitoring of bulk vCA1 terminal activity in vmPFC, NAc, and BA. AAV encoding jRGECO1a was injected into vCA1 and optic fibers were implanted into vmPFC, NAc, and BA. Insets show representative jRGECO1a expressions in vCA1 and projecting areas. Scale bar, 200 μm. **(b)** Aversive conditioning and extinction paradigms. **(c)** Freezing levels to CS^-^ and CS^+^ throughout the experiment (n = 12). **(d)** Mean terminal activity in vmPFC (left), NAc (middle), and BA (right) during the first and last CS^+^ trial in aversive conditioning. **(e)** Mean Z-score peak during the CS (top) and shock period (bottom) of aversive conditioning. CS response was increased at vmPFC and BA terminals of vCA1 during conditioning. Shock response was decreased at vmPFC terminals. **(f)** Mean terminal activity during CS^early^ and CS^late^ of extinction1. **(g)** Mean AUC during the CS (top) and post-CS (bottom) periods. CS-evoked response was decreased in BA terminals during extinction. Post-CS responses were diminished at all terminals.

During aversive conditioning, vCA1 terminals in the vmPFC, NAc, and BA exhibited robust shock-evoked Ca^2+^ responses (**Figure 2d, e**). CS-evoked activity in vmPFC and BA terminals was enhanced during aversive conditioning (**Figure 2d, e**). In extinction, CS responses in BA terminals were significantly higher during CS^early^ compared to CS^late^ (**Figure 2f, g**). The findings indicate that vCA1 transmits signals to BA which dynamically represent associative strength, exhibiting heightened activity during aversive conditioning and a progressive decline in CS-evoked responses throughout extinction. Notably, all vCA1 terminals showed a suppression during the post-CS period in CS^early^ (**Figure 2f, g; Extended Data Fig. 4b**). Post-CS suppressions at all vCA1 terminals diminished in the subsequent session (**Extended Data Fig. 5a, b**), suggesting that post-CS suppressions reflect error of predicted outcome, consistent with single cell imaging (**Figure 1e-g**). Furthermore, all terminals exhibited slow transients to omission responses (Supplementary Table 3), indicating that vCA1 transmits suppression signals, consistent with CS_up_Post_down_ cells observed in single cell imaging. Together, the distinct error signal in vCA1 is broadcast to multiple projection targets.

### vCA1 omission-related signals resist extinction learning

To prove the functional role of post-CS suppression signals, we performed a series of optogenetic experiments targeting vCA1 axon terminals. First, to artificially enhance the inhibitory responses seen during post-CS period (gain-of-function study), we injected AAVs encoding the red-shifted inhibitory opsin, Jaws, or control fluorescent protein into the bilateral vCA1 and implanted optic fibers above either the vmPFC, NAc, or BA (**Figure 3a, b; Extended Data Fig. 6a, b**). During aversive conditioning, where no laser stimulation was delivered, freezing responses did not differ between Jaws and control groups for any of the three terminals. For vCA1-vmPFC terminals, optogenetic inactivation during post-CS period of extinction did not affect freezing responses during CS period of extinction, within-session extinction, but impaired extinction retrieval (**Figure 3c**). For vCA1-NAc terminals, optogenetic inactivation during post-CS period of extinction impaired within-session extinction, but did not alter the extinction retrieval (**Figure 3c**). Likewise, post-CS inactivation of vCA1-BA terminals did not affect within-session extinction but impaired extinction retrieval (**Figure 3c**). Notably, post-CS inhibition of either terminal had no effect on freezing during the post-CS periods of the extinction (**Extended Data Fig. 6c, d**).

**Figure 3.**
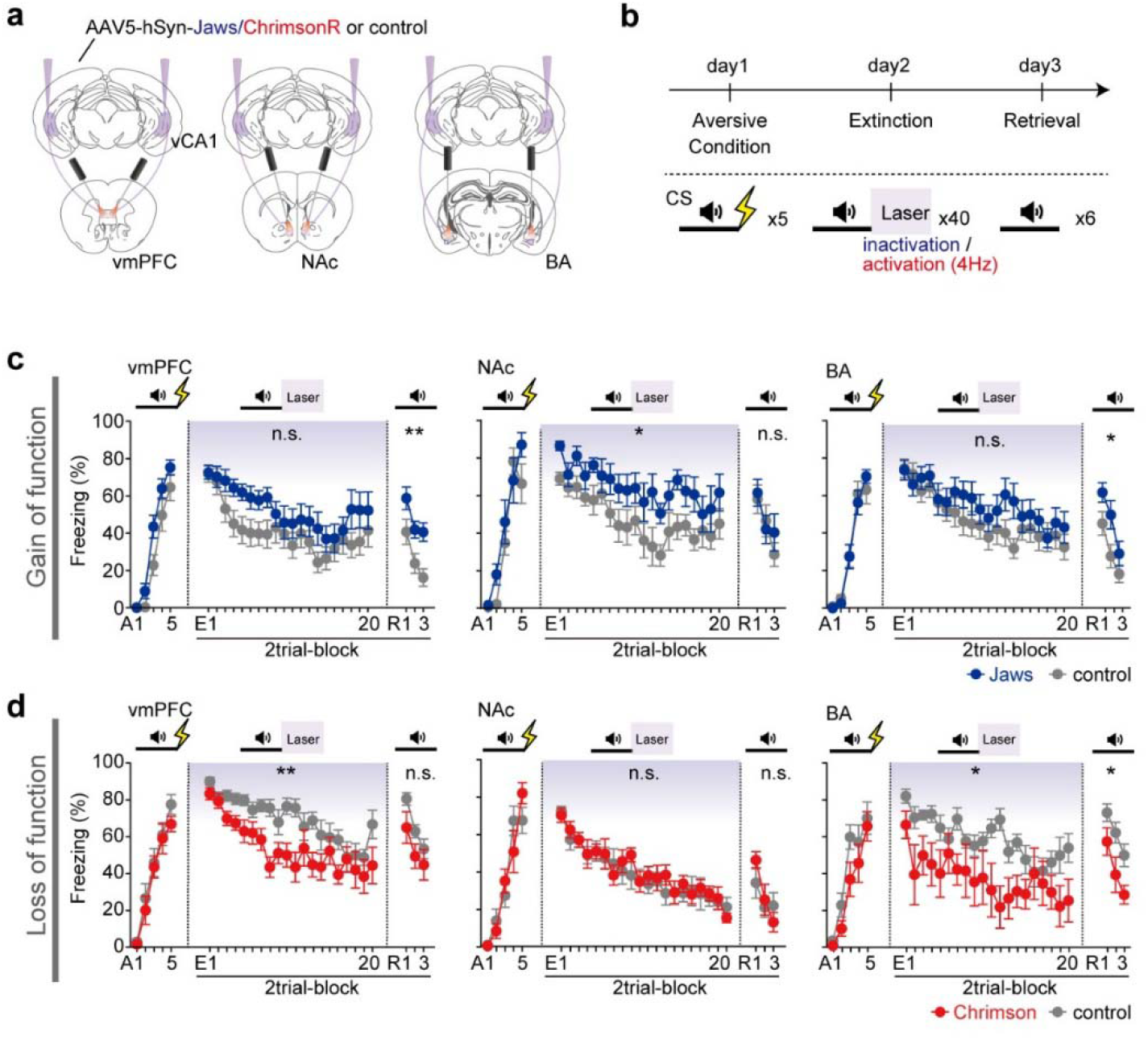
Post-CS activity at vCA1 axon terminals promotes resistance to aversive extinction. **(a)** Terminal manipulation strategy. AAV encoding ChrimsonR, Jaws, or fluorescent protein was injected into vCA1 for optogenetic activation, inhibition, or control, respectively. Optic fibers were bilaterally implanted onto one of vmPFC, NAc, or BA. **(b)** Optogenetic activation or inhibition of terminal activity during the post-CS period of extinction. **(c)** Gain of function study. Freezing during CS in inhibition and control groups. Optogenetic post-CS inhibition of vmPFC terminals (left, n = 12 Jaws, 16 control) and BA terminals (right, n = 14 Jaws, 16 control) impaired long-term extinction. Optogenetic inhibition of NAc (middle, n = 7 Jaws, 8 control) impaired within-session extinction. **(d)** Loss of function study. Freezing during CS in optogenetic activation and control groups in vmPFC (left, n = 8 ChrimsonR, 10 control), NAc (middle, n = 8 ChrimsonR, 11 control), and BA (right, n = 7 ChrimsonR, 11 control). Optogenetic post-CS activation of vmPFC terminals facilitated within-session extinction. Optogenetic post-CS activation of BA terminals facilitated within-session extinction and extinction retrieval. A, aversive conditioning; E, extinction; R, retrieval. n.s., not significant; \**P* < 0.05; \*\**P* < 0.01; For statistical details, see Supplementary Table 1.

Next, we optogenetically activated each projection pathway during extinction (loss-of-function study) using Chrimson, a red-shifted excitatory opsin, expressing mice (**Figure 3a, b; Extended Data Fig. 7a, b**). During aversive conditioning, freezing responses did not differ between Chrimson and control groups for any of the three terminals. For vCA1-vmPFC, optogenetic activation during post-CS period of extinction facilitated within-session extinction without affecting the extinction retrieval (**Figure 3d**). Unlike inactivation, post-CS activation of vCA1-NAc terminal did not show a significant impact on freezing during extinction and retrieval (**Figure 3d**). For vCA1-BA, optogenetic activation during post-CS period of extinction facilitated within-session extinction as well as extinction memory retrieval (**Figure 3d**). Like optogenetic inhibition, post-CS activation of either terminal had no effect on freezing during the post-CS periods of the extinction (**Extended Data Fig. 7c, d**). These data demonstrate that the vCA1 shock-omission signals act as resistance to change signals that suppress extinction learning, with subtle differences across the projection target, standing in contrast to dopamine prediction error signal that promotes extinction of aversive memory ^25,34^.

### vCA1 computes latent state inference errors

Distinct from the canonical prediction error frameworks ^1-7^, our findings revealed that vCA1 encodes a non-canonical function of error signals in response to expected outcome omission, which restrains new learning. According to the latent cause model, the hippocampus is proposed to infer the latent cause underlying an observation through Bayesian inference ^12^. Within this framework, we hypothesized that the behavior maintaining function may be driven by a belief state that maintains prior task representations and resists new learning, as described in a Bayesian inference theory ^35^. Because the relationship between sensory observations and their underlying latent causes cannot be uniquely determined from observations, inference over latent states necessarily relies on internal belief representations. We therefore used a Partially Observable Markov Decision Process (POMDP) within an active inference framework to estimate the animal’s belief about aversive and safety contexts, which correspond to latent task-relevant structures during aversive conditions and extinction (**Figure 4a**). This framework allowed us to calculate both discrete belief transitions and the moment-by-moment belief probabilities. We designed the POMDP such that each belief of the POMDP corresponds to a distinct behavior strategy. For instance, in an “aversive context” state, animals exhibited high likelihood of defensive responses, whereas in a “safe context” state, animals exhibited low likelihood of responses. Finally, each inference error is defined as the time derivative of the corresponding inference probability.

**Figure 4.**
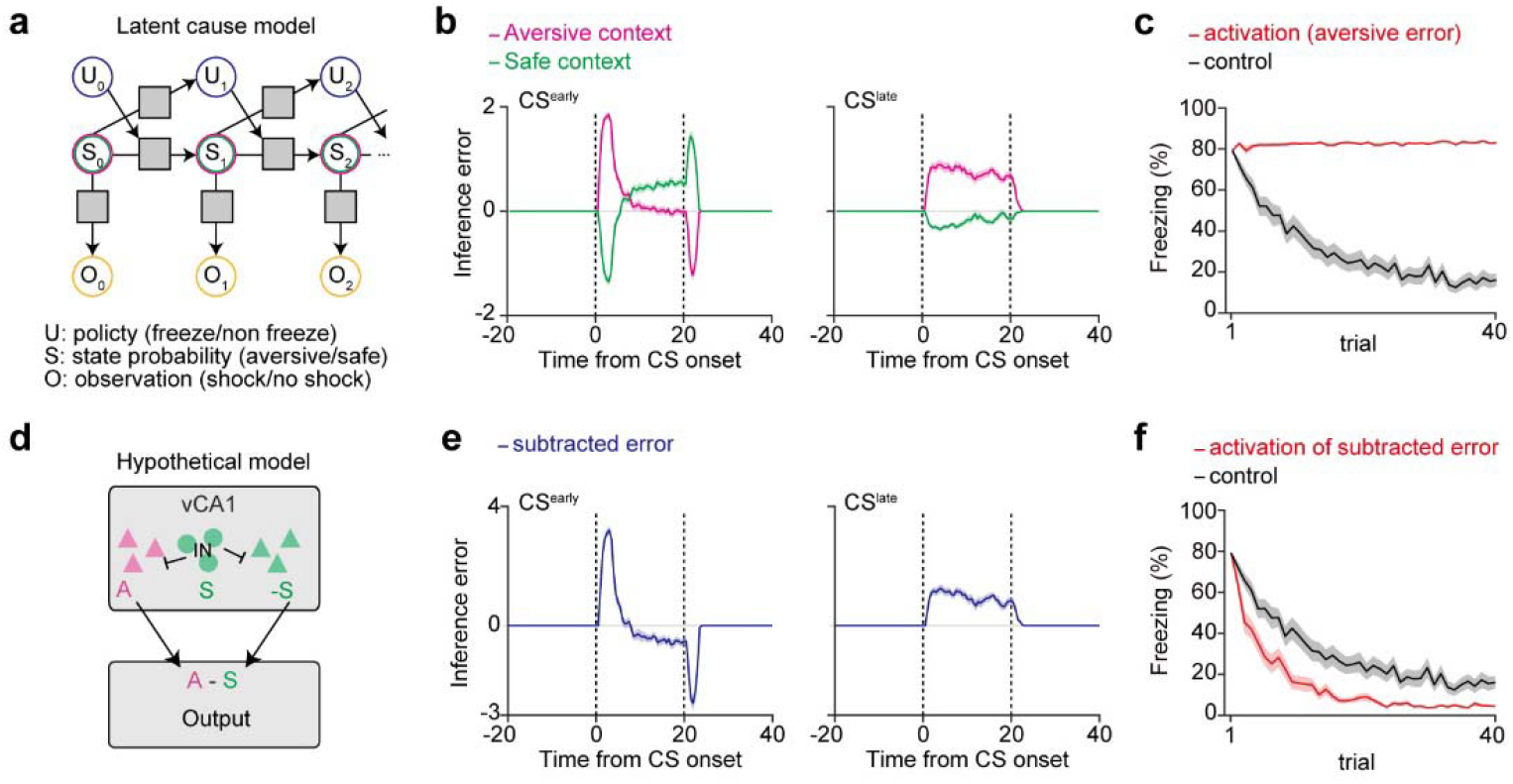
vCA1 neurons represent inference errors during aversive extinction in the simulation model. **(a)** Schematic of the latent cause model. Gray squares represent transition probabilities (between S_t_ and S_t+1_), policies (between S_t_ and U_t+1_), and likelihood (between S_t_ and O_t_). **(b)** Inference errors of aversive and safe context during CS^early^ and CS^late^ of extinction session. **(c)** Loss of function study. Simulated freezing levels during the CS period with manipulation of aversive inference error. Freezing levels increased when aversive inference error signals were attenuated during the post-CS period. **(d)** A hypothetical circuit model illustrating how vCA1 integrates inference errors between aversive and safe context. Interneuron (IN) signals safe context, which is in turn converted to negative signal to output regions. **(e)** Inference errors in the subtraction model during CS^early^ and CS^late^ of extinction session. **(f)** Loss of function study. Simulated freezing levels during the CS period with manipulation of subtracted inference error. Freezing levels decreased when subtracted inference error signals were attenuated during the post-CS period.

In this framework, we found that inference errors for the aversive context decreased during the post-CS period, whereas inference errors for safety contexts increased during the post-CS period (**Figure 4b**), in which inference error dynamics of aversive context closely follows vCA1 terminal activity (**Figure 2f**). In the simulated model, artificially reducing the negative component of the aversive inference error during the post-CS period toward zero (loss-of-function study) led to an extinction deficit (**Figure 4c**). Notably, this result differs from the experimental observation that optogenetic disruption of post-CS signals facilitated extinction learning (**Figure 3d**), suggesting that vCA1 terminal activities reflect more than aversive contextual belief state. Together, these results imply vCA1 signals integrate additional computational component beyond aversive inference.

Because different forms of aversive conditioning and extinction recruit largely distinct populations of dorsal hippocampal neurons with a small overlap ^36-38^, we hypothesized that vCA1 contains neuronal subsets representing inference errors for aversive and safe context and that vCA1 sends a composite signal reflecting both aversive and safe inferences to downstream regions. Given that vCA1 interneurons (INs) have been implicated in facilitating appetitive and aversive extinction ^39,40^, we designed a subtraction model in which subsets of projection neurons and INs are aversive and safe context inference nodes, respectively. Through the local connection, aversive neurons and/or distinct projecting neurons receive inhibitory information of IN encoding inference errors for safety context (**Figure 4d**). This model reproduced the temporal features observed in vCA1 terminals during both early and late phase of extinction learning (**Figure 4e**). When we artificially attenuated the negative component of inference error signals at the post-CS period in the subtraction model (loss-of-function study), extinction learning was enhanced (**Figure 4f**), consistent with our biological observation. These results support the idea that vCA1 reflects a hybrid of inference errors for both aversive and safe context.

### Resistance to change signals are prominent during expectation-guided behavior

According to psychological theories of habit formation, repeated actions serve to consolidate and maintain the underlying beliefs that generated them ^41^. Likewise, when freezing is directed by a strong belief of aversive state, freezing could provide feedback that stabilizes the current belief state. In an active inference framework, actions influence perceived sensory inputs, which in turn update beliefs about hidden states. To investigate behavior-guided signals, we analyzed inference errors during post-CS freezing periods in our simulation model. We found that inference errors in a subtraction model decreased during post-CS freezing epochs (**Figure 5a-d**), indicating freezing-induced inhibition of belief updating. To examine how biological signals correspond to these computational predictions, we then analyzed post-CS freezing-triggered activity from single cell and fiber photometry recordings. In single cell recording, all Post_down_ types including CS_up_Post_down_ cells, but not Post_up_ cells, decreased activity in a post-CS freezing-specific manner (**Figure 5e-h; Extended Data Fig. 8a-e**). Specifically, the post-CS Ca^2+^ signal decreased around the onset of freezing and increased around the offset (**Figure 5g, h; Extended Data Fig. 8a-e**). In line with these Post_down_ cells, fiber photometry data revealed that vCA1-vmPFC, - NAc, and -BA terminals also showed freezing-specific reductions in activity during post-CS period throughout the extinction session (**Figure 5i-k; Extended Data Fig. 9a-c**). Furthermore, the post-CS AUC was negatively correlated with the post-CS freezing level (**Extended Data Fig. 9d**). These freezing-specific reductions could be due to speed cells in the hippocampus, where the firing rates positively correlate with locomotion speed in the dorsal CA1 ^42-44^. To assess the effect of locomotion, we analyzed immobility onset or offset during CS^-^ trials in aversive conditioning, where no shock was delivered. Across all regions, no differences in terminal activity were observed between the CS^-^ immobility and non-immobility periods in aversive conditioning, indicating that locomotion effect would be negligible in all terminals (**Extended Data Fig. 9e, f**). Together, vCA1 neurons represent robust belief-maintenance signals particularly during freezing response, which is thought to modulate belief updating in active inference model.

**Figure 5.**
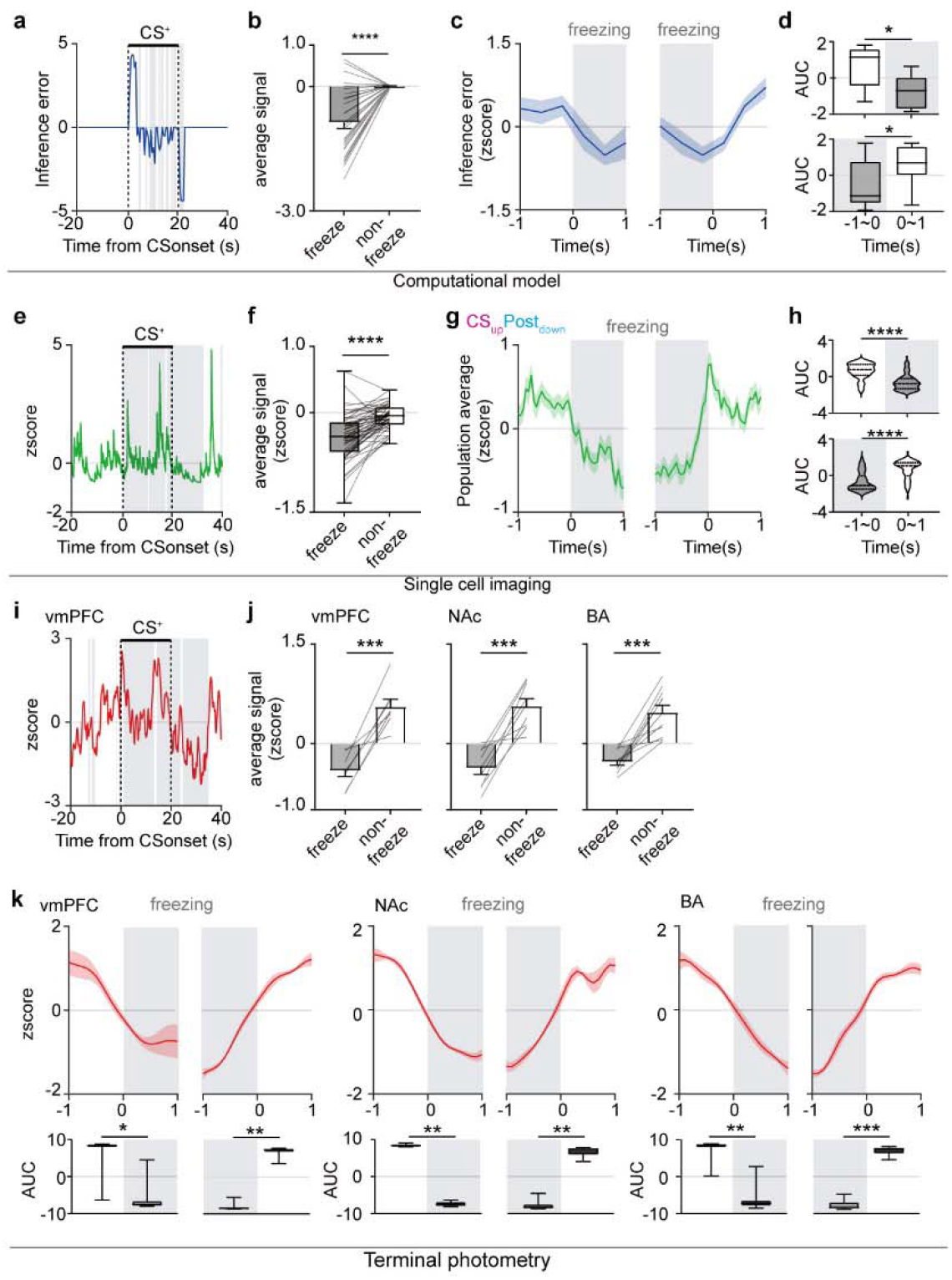
Behavior-dependent belief-maintenance signals in vCA1. **(a)** Representative trace of subtracted error from computational model. The gray shaded area indicates freezing period. **(b)** Mean subtracted inference error signals during post-CS freezing or non-freezing period without temporal constraints in computational model. Subtracted inference errors during freezing period were lower than those during non-freezing period. **(c)** Subtracted inference error signals in the simulated model aligned to the onset (left) and offset (right) of post-CS freezing. Freezing epochs were extracted only when no behavior transition occurred within 1 s before or after the epoch. The gray shaded area indicates freezing period. **(d)** Mean AUC decreases during freezing period for both alignments. White bars denote non-freezing and gray bars indicate freezing periods. Subtracted inference error decreased at post-CS freezing onset and increased at post-CS freezing offset. **(e)** Representative trace from CS_up_Post_down_ cell from single cell calcium imaging during extinction. The gray shaded area indicates freezing period. **(f)** Mean signal of CS_up_Post_down_ cells during post-CS freezing or non-freezing period without temporal constraints during extinction. Responses during freezing period were lower than those during non-freezing period in CS_up_Post_down_ cells. **(g)** Population activity of CS_up_Post_down_ cells from single cell calcium imaging, aligned to post-CS freezing onset (left) and offset (right). Freezing epochs were extracted only when no behavior transition occurred within 1 s before or after the epoch. **(h)** Violin plots of individual cell AUC during 1sec of freezing and non-freezing epochs. CS_up_Post_down_ responses decreased at freezing onset and increased at freezing offset. **(i)** Representative photometry trace from vCA1-vmPFC terminals during extinction. **(j)** Mean terminal responses during post-CS freezing or non-freezing period without temporal constraints during extinction. Responses during freezing period were lower than those during non-freezing period. **(k)** Mean z-score responses aligned to post-CS freezing onset and offset during extinction in all vCA1 terminals. Freezing epochs were extracted only when no behavior transition occurred within 1 s before or after the epoch. Insets show mean AUC decreases during freezing period for both alignments. vCA1 terminal responses decreased at freezing onset and increased at freezing offset.

### Behavior-specific vCA1 signals prevent extinction learning

Finally, we assessed whether the freezing-specificity of vCA1 belief-maintenance signals is functionally important. We employed a closed-loop optogenetic approach to manipulate the post-CS terminal activity of vCA1 neurons time-locked to freezing during aversive extinction (**Figure 6a**). Mice were injected with AAV5-hSyn-Jaws or control fluorescent protein followed by bilateral optic fiber implantations above one of vmPFC, NAc, or BA. We used a real-time pose estimation system to detect freezing ^45^, which triggered red laser delivery either during post-CS freezing or non-freezing periods (**Figure 6a**). During aversive conditioning, in which no laser stimulation was delivered, freezing responses did not differ between the Jaws and control groups for any of the three terminals (**Figure 6b, c**). Post-CS inhibition of the vCA1-vmPFC terminals during extinction impaired extinction retrieval, but not within-session extinction, in the freezing-triggered group (**Figure 6b**). For vCA1-NAc terminals, freezing-specific inhibition during the post-CS period of extinction impaired within-session extinction without affecting extinction retrieval (**Figure 6b**). These effects were specific to the freezing-triggered condition, as neither manipulation in the non-freezing-triggered group altered within-session extinction nor extinction retrieval (**Figure 6c**). For vCA1-BA terminals, post-CS inhibition did not affect within-session extinction or extinction retrieval in either the freezing or non-freezing condition (**Figure 6b, c**), indicating pathway-specific and behavior-dependent roles of vCA1 signals. In addition, neither freezing nor non-freezing specific inhibition affected freezing levels during the post-CS periods throughout the extinction session in any of the terminals (**Extended Data Fig. 10a-d**). Notably, the duration of inhibition for the freezing-triggered group was shorter than that for the non-freezing-triggered group (**Extended Data Fig. 10e**), ruling out the possibility that extinction was impaired only in the freezing-triggered group due to a longer inhibition duration. Together, freezing-locked post-CS activity in vCA1 terminals conveys belief-maintenance signals that causally constrain extinction.

**Figure 6.**
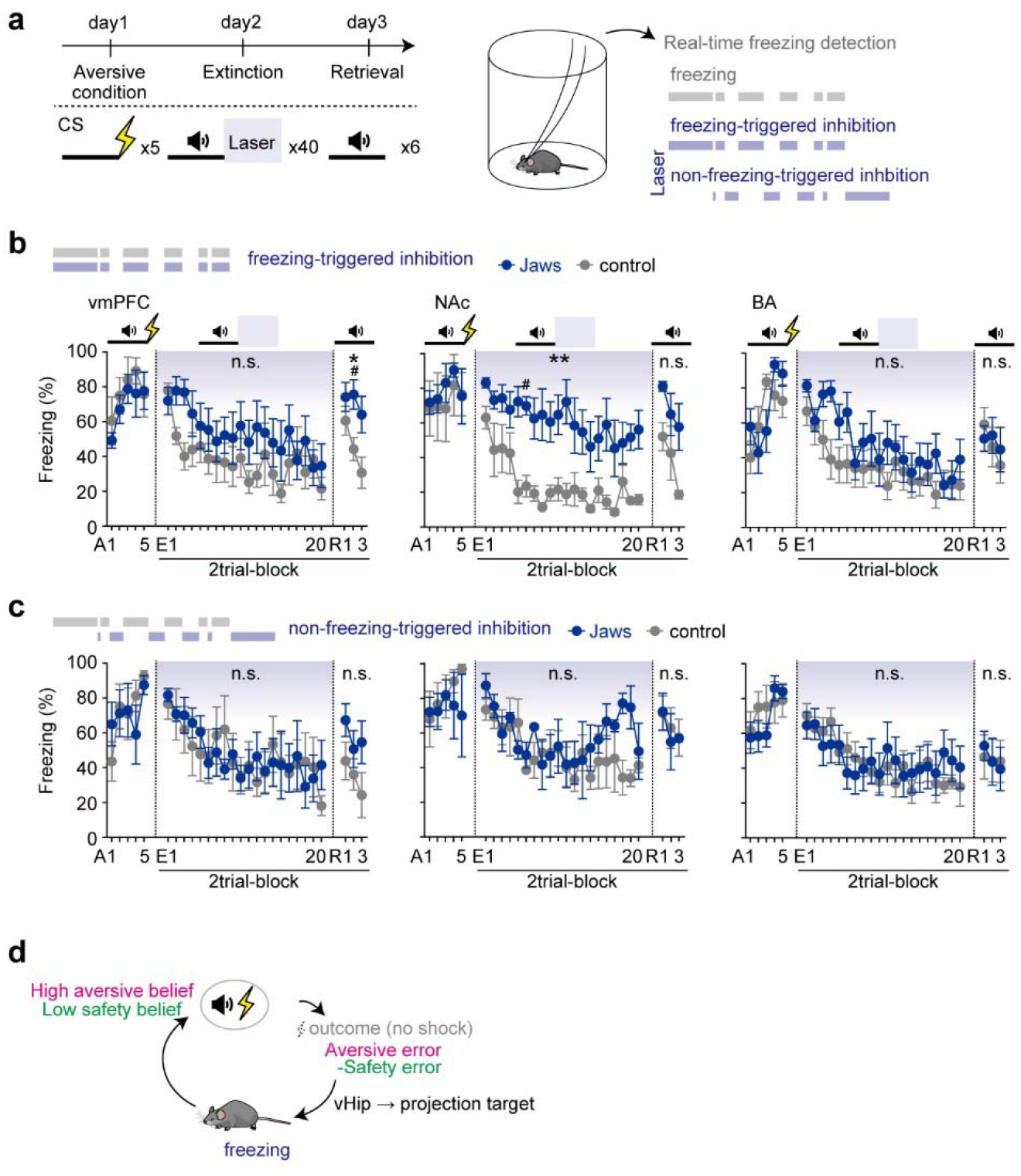
Freezing-dependent vCA1 signals resist aversive extinction. **(a)** Closed-loop optogenetic inhibition during the post-CS period. **(b, c)** Gain of function. Freezing levels during CS in freezing-triggered (a) or non-freezing-triggered (b) groups. Freezing-triggered inhibition of vmPFC and NAc terminal activity impaired long-term extinction memory and within-session extinction. No statistically significant effects in non-freezing triggered groups. A, aversive conditioning; E, extinction; R, retrieval. **(d)** Model of vCA1 inference error signals. n.s., statistically not significant; \**P* < 0.05; \*\**P* < 0.01; \*\*\**P* < 0.001; \*\*\*\**P* < 0.0001; For statistical details, see Supplementary Table 1.

## DISCUSSION

We show that vCA1 is inhibited at the omission of predicted outcome, especially when mice show expectation-guided behavior. Although prediction error signals have been conceptualized as teaching signals that drive associative learning by updating expectations to align with outcomes ^1,46^, our findings reveal a qualitatively different role of error coding in the hippocampal circuit. The inhibitory response we observe in vCA1 arises from a mismatch between inferred contextual states and actual sensory inputs, but instead of promoting updating, it acts to resist extinction learning and maintain an aversive state. Thus, in contrast to reinforcement learning frameworks, the vCA1 error signal operates in opposition to new learning. Our simulation model formalizes this computation as an inference error signal that integrates sensory inputs with inferred contextual states. Under this formulation, resistance to change reflect precision weighting of prior belief rather than failure of new learning (Supplementary Information). These results indicate that vCA1 encodes a form of error signal linked to internal expectations that stabilizes aversive memory representations and suppress premature updating (**Figure 6d**).

Resistance to switch to a new internal model can be adaptive in at least two ways. First, suppressing immediate updating allows the system to preserve previously acquired aversive memories, which may remain valuable for survival if aversive contingencies reoccur ^47^. Second, maintaining prior state inference allows context-dependent memory retrieval such as fear renewal in distinct environments ^48-51^. Thus, the resistance to extinction in vCA1 may serve as an ecological function to prevent premature overwriting of aversive memories while permitting flexible memory reactivation depending on environment. In parallel, other neural systems such as dopamine pathways promote extinction ^25,34^, suggesting that opposing processes operate concurrently to balance updating internal models.

Although terminal photometry experiments revealed enhanced CS responses and reduced post-CS responses during CS^early^, single-cell imaging uncovered diverse patterns of CS and post-CS responses across individual neurons. This heterogeneity suggests that population-level signals arise from selective recruitment of functionally distinct subpopulations, including neurons with CS_up_Post_down_ dynamics that may disproportionately contribute to terminal signals.

Mechanistically, inhibitory circuit dynamics are likely to contribute to post-CS suppression. Somatostatin interneurons have been implicated in extinction and exhibit activity linked to freezing behavior^40^. In our data, a subset of neurons displayed Post_up_ responses associated with freezing, raising the possibility that inhibitory populations contribute to stabilizing inferred states or gating safety-related signals. Although direct cell-type-specific evidence is required, these findings point to inhibitory control as a key mechanism regulating state maintenance in vCA1.

While hippocampal principal neurons exhibit rapid plasticity in response to excitatory input^52^, interneurons such as somatostatin- or parvalbumin-expressing cells may provide counteracting inhibition that stabilizes existing ensembles and limits immediate reorganization^39,40,53^. Notably, post-CS suppression occurs within a temporal window critical for plasticity induction, suggesting that inhibitory dynamics may act to gate learning at the moment when updating would otherwise occur. Consistent with this idea, CS_up_Post_down_ neurons exhibit stronger CS responses than CS_up_Post_up_ neurons during late extinction (**Extended Data Fig. 1g**)., and computational modeling indicates that inhibition is necessary for maintaining aversive state representations (**Figure 4f**). Together, these findings suggest that post-CS inhibition biases the vCA1 circuit toward maintaining inferred states rather than updating them, positioning the hippocampus as a circuit that regulates whether learning should occur.

## Methods

### Animals

All experimental procedures were approved by the Animal Care Committee of the University of Tokyo and the National Institute of Advanced Industrial Science and Technology. Male C57BL/6J mice, aged 8-12 weeks old at the time of surgery, were used for all experiments. The mice were group-housed three to four in a cage prior to surgery and subsequently transferred to individual housing. All mice were maintained in a ventilated and temperature-controlled environment (temperature, 23 ± 2 °C, humidity, 50 ± 10 %), under a 12-hour light/dark cycle (lights on 8 a.m. to 8 p.m.), with *ad libitum* access to food and water. All behavioral experiments were conducted during the light cycle.

### Viruses

The following adeno-associated viruses (AAV), along with the respective suppliers, injection titers, and injection volumes, were used in this study: AAV9-hSyn-RiboL1-jGCaMP8s (International Research Center for Neurointelligence; IRCN, 2.41 × 10^13^ vg/ml, 500 nL per injection) and AAV5-hSyn-jGCaMP8s (IRCN, 2.69 × 10^13^ vg/ml, 500 nL per injection) for the miniature microscope imaging, AAV9-hSyn-NES-jRGECO1a (Addgene, 1.8 × 10^13^ vg/ml, 150 nL per injection) for the multi-fiber photometry imaging, AAV5-hSyn-Chrimson-TdT (IRCN, 1.0 × 10^13^ vg/ml, 150 nL per injection) for optogenetic activation, AAV5-hSyn-Jaws-KGC-GFP-ER2 (IRCN, 9.7 × 10^12^ vg/ml, or Addgene, 2.0×10^13^ vg/ml, 150 nL per injection) for optogenetic inhibition, AAV5-hSyn-EYFP (UNC, 6.6 × 10^12^ vg/ml, 150 nL per injection) or AAV5-hSyn-EGFP (IRCN, 1.0 × 10^13^ vg/ml, 150 nL per injection) as a control for optogenetic activation and inhibition.

### Surgical procedures

The mice were anesthetized with isoflurane (Viatris, VTRS, 3% for induction, 1-2% for maintenance) at a constant flow rate with an anesthesia pump (RWD) and secured in a stereotaxic frame (RWD or Kopf Instruments). Throughout the surgical procedure, the mice were placed on a heating pad to ensure that their body temperature was maintained at an optimal level. Moreover, a Vaseline solution (KENEI Pharmaceutical) was applied to prevent the eyes from desiccating. Prior to incision of the skin, a combination of antibiotics (Bayer, Baytril 2.5% injection solution for dogs and cats) and analgesics (Boehringer Ingelheim, Metacam 2% solution for injection) was administered subcutaneously in a volume of 0.1 ml. A gel (Jex, Projelly) was applied to the cranial opening to prevent desiccation of the brain surface. The virus was injected into the target area using a pulled-glass capillary (Drummond, Nanoject Glass Capillaries) and a microinjector (Drummond, Nanoject III) at a flow rate of 1 nL per second. Subsequently, the microinjector was left in place for 10 minutes to prevent backflow of the injected virus. The GRIN lens and optic fibers were affixed to the skull with UV Adhesive (Bondic, BD-CRJ), Super-Bond (Sun Medical, Super Bond C & B Set), and acrylic dental cement (Yamahachi Dental MFG., Re-fine Bright).

For the miniature microscope imaging, AAV9-hSyn-RiboL1-jGCaMP8s or AAV5-hSyn-jGCaMP8s was unilaterally injected into the vCA1 (anteroposterior (AP) -3.30 mm, mediolateral (ML) 3.50 mm (right), dorsoventral (DV), -3.70 and -3.80 mm or AP -3.30 mm, ML 3.75 mm (right), DV -4.20 and -4.30 mm from bregma). Following the injection, a GRIN lens (GoFoton, CLES050GFT100, diameter: 0.5 mm, length: 8.09 mm) was unilaterally implanted into the vCA1 (AP -3.30 mm, ML 3.50 mm (right), DV -3.75 mm or AP -3.30 mm, ML 3.75 mm (right), DV -4.25 mm from bregma). Three to 12 weeks following the surgical procedure, mice were placed in a custom-made head-fix apparatus, where a baseplate (Open Ephys, Miniscope V4 Base Plate) was affixed to the dental cement at the focal point.

For the multi-fiber photometry imaging, AAV9-hSyn-NES-jRGECO1a was unilaterally injected into the vCA1 (AP -3.16 and -3.52 mm, ML 3.50 mm (right), DV -3.60 and -3.90 mm from bregma). Optic fibers (Newdoon, ferrule diameter: φ1.25 mm, fiber core: 200μm, fiber length: 6.0 mm) were then unilaterally implanted into the vmPFC (AP 1.80 mm, ML 1.60 mm (right), DV -2.00 mm at an angle of 30° in the ML direction from bregma), NAc (AP 1.20 mm, ML 0.58 mm (right), DV: -4.70 mm from bregma), and BA (AP -2.06 mm, ML 3.30 mm (right), DV -4.60mm or -4.70mm from bregma).

For optogenetic manipulation experiments, AAV5-hSyn-Chrimson-TdT, AAV5-hSyn-Jaws-KGC-GFP-ER2, AAV5-hSyn-EYFP, or AAV5-hSyn-EGFP was bilaterally injected into the vCA1 (AP -3.16 and -3.52 mm, ML ±3.50 mm, DV -3.60 and -3.90 mm from bregma) and optic fibers were bilaterally implanted in the vmPFC (AP 1.70 mm, ML 1.68 mm, DV - 2.50mm at an angle of 30° in the ML direction from bregma), NAc (AP 1.20 mm, ML 1.78 mm, DV -4.34 mm at an angle of 15° in the ML direction from bregma), or BA (AP -2.06 mm, ML 3.30 mm, DV -4.70 mm from bregma).

### Aversive Conditioning and Extinction in Calcium Imaging Experiments

Behavioral experiments were conducted 4-12 weeks after the surgery for the miniature microscope imaging and 6 weeks after the surgery for the multi-fiber photometry imaging, allowing for sufficient recovery from surgery and virus expression. The animals were handled weekly until the behavioral experiments were conducted. All behavioral experiments were conducted in a soundproof chamber with stimuli controlled by data acquisition system (O’hara Co., Ltd.) or pyControl (OpenEphys). To avoid the potential influence of circadian rhythms, each mouse was subjected to a series of behavioral experiments at approximately the same time each day. In aversive conditioning and extinction paradigm, mice received an auditory conditioned stimulus (CS) and/or electrical foot shock unconditioned stimulus (US). CSs were pure tones (4 kHz and 10kHz, 20s, counterbalanced); CS^+^ was followed by US (2s, 0.4 mA) immediately after CS offset, whereas CS-was not. On Day1, mice were presented with 30 CS^-^ within their homecage for habituation. On Day2, mice were conditioned with 5 pairings of CS^+^ and US after initial 5 CS^-^ presentations. On Day3 (and Day4 for the multi-fiber photometry imaging), mice received 5 CS^-^ followed by 40 CS^+^ alone presentations in distinct context from aversive condition. On Day4 (or Day5 for the multi-fiber photometry imaging), mice were exposed to 5 CS^-^ followed by 10 CS^+^ alone presentation in extinction context. All trials were separated with variable intertrial intervals (100 s on average for conditioning and 70 sec on average for extinction and retrieval).

All behavioral sessions were video-recorded with an overhead camera (Basler, acA1920-120um or ARGO, DMK22BUC03) at 20 frames per second. DeepLabCut was used to extract several body points ^54^. The accuracy of the labeling was verified by visualizing the videos with the labeling markers using an existing program within DeepLabCut. Freezing periods were defined as periods in which the positional changes of these body points between consecutive frames were below a threshold for at least 1 sec. Then, freezing periods were validated through a subsequent manual inspection. Freezing periods were identified when mice exhibited no discernible movement apart from respiration. In miniscope experiments, freezing periods were determined solely by manual inspection. Based on the synchronized timestamps, freezing periods were aligned to imaging frames for further analyses.

### Reward Training and Omission in Calcium Imaging Experiments

Following aversive extinction, the subset of animals was subjected to reward paradigm. Mice were habituated to 5% sucrose solution one week before the experiment. They were water-deprived throughout the reward experiment, with 1h of daily water access to maintain body weight at 90% of the baseline level. Mice were head-fixed and placed close to a lick spout in sound-proof chamber during all sessions. The US was a 5% sucrose (3ul), delivered immediately after CS offset. On Day 0, water-restricted mice were subjected to 2 habituation sessions for an experimental setup separated by at least 2 hours. Mice were presented with sucrose solution for 120 and 60 times in the first and second session, respectively. On Day1 and 2, mice received 120 CS^r^ (7.5 kHz tone, 1s, 74dB), each immediately followed by sucrose delivery. On Day3, mice were presented with CS^r^ with or without sucrose delivery at a 4:1 ratio (omission randomized). All trials were separated with variable intertrial intervals (25 sec on average). Licking was detected with a touch sensor equipped with a spout (Bit Trade One, ADFCS01). Lick counts were visualized with 250-millisecond time bins, averaged across sucrose or omission trials using custom written code in Matlab.

### Miniature Microscope Imaging

Imaging was conducted using a miniature microscope (OpenEphys, Miniscope V4.4). Mice were habituated to the miniature microscope attachment procedure prior to behavioral experiments. Images were acquired at a frame rate of 20 Hz with an LED power of 55-85 % (0.89-2.20 mW at the objective, 470 nm) depending on jGCaMP8s expression, analog gain of 2, and with 608 × 608 pixels. To ensure millisecond precision, the timestamps of the imaging frames and experimental TTLs were all collected for alignment using the pyControl breakout board (OpenEphys).

### Analysis of Miniature Microscope Imaging Data

#### Pre-processing

A series of preprocessing steps was initially applied to the recorded movies, which includes cropping, band-pass spatial filters (Low cut-off, 0.005 pixel^-1^, High cut-off, 0.500 pixel^-1^), and motion correction (Inscopix Inc.). For cell identification, Constrained Non-negative Matrix Factorization for endoscopic data (CNMFe) was applied to the preprocessed data. Briefly, a pixel whose temporal correlation with the 8 neighbor pixels that surround it is high (> 0.7) and peak-to-noise ratio is high (> 6) is selected as a seed pixel, and the seed pixel is considered to be a potential cell. Based on the pre-defined cell diameter (10-12 pixels), a basic cell footprint around the seed pixel and its trace are extracted. Then, these footprint and traces serve as the initial guess to the CNMF optimization to extract spatial and temporal features of each cell from the movies. Automatically identified cells and their traces were manually confirmed, and those considered not to be cells were excluded from the data. Neurons detected on aversive conditioning and extinction were aligned according to their region of interest (ROI) longitudinally using a cell registration algorithm. For analysis of data from aversive conditioning, cells which were detected both in aversive conditioning and extinction were used, while for analysis of data from extinction, cells detected in each experiment were used. The extracted signals were further processed using custom-written codes on MATLAB.

#### Supervised Clustering

For the classification of functional types, averaged signals during CS, US, and post-CS signals were compared to pre-CS baseline signals. Specifically, 0-5s period from CS onset and equivalent baseline period were compared for CS classes, 0-2s period from US onset and equivalent baseline period were compared for US classes in conditioning, and post-CS 0-20s period and equivalent baseline period were used for post-CS classes in extinction. Trial-binned values were used for CS and post-CS analyses, while all single trial values during US were used for US analyses. Responsive cells were identified if their time-binned values (1sec) during the stimulus were significantly increased or decreased. Normality of the data was not assumed, and all tests were nonparametric tests (Wilcoxon signed-rank test).

During aversive extinction, the peristimulus time histogram (PSTH) aligned to CS onset of individual cell was calculated by averaging block of 5 trial bins triggered by CS onset and converting trial-binned values to *Z*-scores using the formula *Z*_*i*_ = (*S*_*i*_ - μ)/σ , where μ and σ are the mean and s.d. of baseline frames at each trial-bin, respectively. To assess the trial difference during extinction, the area under curve (AUC) during CS or post-CS period were compared across the CS-, CS^early^, and CS^late^ in each cell type. For aversive conditioning, the PSTH aligned to CS onset of individual cell was calculated by single trial values triggered by CS onset and converting values to *Z*-scores. Population activity of each functional cell type was calculated based on *Z*-scores. To see the trial difference during conditioning, the AUC during CS or shock period were compared between the first and last trials in each cell type.

#### SVM

For constructing time-resolved support vector machine (SVM), we used a linear SVM. For each trial, feature vectors were constructed by concatenating the mean values of individual neurons computed within a 5-s time window. Decoding was performed using a dataset consisting of 10 trials (5 CS- and 5 CS^early^ trials) at each time point with stratified 5-fold cross-validation. Chance-level performance was estimated by training the classifier with randomly shuffled trial labels (100 iterations). Statistical significance was evaluated by comparing the observed accuracy with the null distribution derived from shuffled labels. To assess how functional cell types contribute to classifier, the average weight of each cell was calculated and compared across functional cell types. Normality of the data was not assumed, and all tests were nonparametric tests.

#### Freezing-triggered Analysis

Calcium responses from each mouse were aligned to binarized behavioral data (freezing or not freezing) and triggered at the onset or offset of freezing epochs during the post-CS period across the entire session. Signals within 1-second window before and after each event were extracted. Epochs were excluded if another freezing event occurred within the time window. Subsequently, the extracted signals were averaged and z-score normalized. Normality of the data was not assumed, and all tests were nonparametric tests (Wilcoxon signed-rank test).

### Multi-fiber Photometry Imaging

To record calcium signals at axon terminals from three distinct brain regions simultaneously, multi-fiber photometry recordings were conducted using the FP3002 system (Neurophotometrics), controlled by Bonsai, as described elsewhere. The 560 nm, 470 nm, and 415 nm (isosbestic) LEDs in the FP3002 system were calibrated to deliver 100 µW, 50 µW, and 50 µW of power at the tip of the fiber, with recordings taken at a 10 Hz sampling rate for each wavelength.

### Analysis of Multi-fiber Photometry

#### Pre-processing

Fiber photometry data were processed for normalization, fitting, and sorting using a custom written program in MATLAB (Mathworks). Motion correction was performed by adjusting the fluorescence of jRGECO1a measured at 560 nm with data from the 415 nm reference point.

#### PSTH

During aversive conditioning, PSTH aligned to CS onset was calculated by individual trial triggered by CS onset. AUC during CS (0-10sec after CS onset) and shock (0-2sec after shock onset) responses were quantified and compared to pre-CS signals. In addition, CS^+^ and shock responses in the first and last trial of were also assessed to learning-induced changes.

For CS responses, 0-10s period from CS onset was used. For shock responses, peaks of shock period were used for conditioning. For post-CS responses during extinction, the AUCs of 0-20 sec after CS offset were used. During aversive extinction and its retrieval, PSTH aligned to CS onset was constructed by averaging blocks of 5 trials. Similar to aversive conditioning, AUC during CS (0-10sec after CS onset) and post-CS (0-20sec after CS offset) responses were quantified and compared.

#### Freezing-triggered analysis

Calcium responses from each mouse were aligned to binarized behavioral data (freezing or not freezing) and triggered at the onset or offset of freezing epochs during the post-CS period across the entire session. Signals within 1-second window before and after each event were extracted. Epochs were excluded if another freezing event occurred within the time window. Subsequently, the extracted signals were averaged and z-score normalized.

### Optogenetic Manipulation

#### Post-CS Manipulation

Optical fiber cables (Thorlabs, BFYL2LF01) were connected to optic fibers to deliver laser bilaterally. For optogenetic activation, a laser at a wavelength of 635 nm (square wave, 4 Hz, 10 ms duration, 7-8 mW) was irradiated during the post-CS period (20 s following CS offset) of the extinction session. As 4 Hz, rather than 20 Hz, has been demonstrated to be effective for CA1 activation ^55^, we adopted a 4 Hz stimulation protocol. For optogenetic inhibition, a continuous laser at a wavelength of 635 nm (15 mW) was irradiated during 20s after CS offset in extinction.

#### Closed-loop manipulation

To obtain a balanced amount of freezing and non-freezing time during the post-CS period, a subset of mice in the photoinhibition group were reconditioned and subsequently subjected to closed-loop manipulation. For online freezing detection, the automatic tracing of real-time body parts was conducted using the DeepLabCut network, which had been already trained with other datasets as previously described. The absolute changes in X and Y coordinates of each body part were summed using Bonsai-DLC ^45^. Subsequently, the values were low-pass filtered with a cutoff frequency of 1 Hz. The freezing was defined as a condition in which the sum of the XY changes of at least one body part fell below the pre-defined threshold for a duration of 0.5 seconds or more. For the freezing-triggered group, the continuous laser at a wavelength of 635 nm (15 mW) was triggered when algorithm detected freezing during the post-CS period (20s after CS offset). This laser illumination was terminated immediately when the algorithm determined non-freezing period or when post-CS period ended. In contrast, for the non-freezing-triggered group, the laser was switched on and off under the opposite conditions.

### Histology

Following behavioral experiments, a histological verification of virus expression and implant placement was conducted. The mice were deeply anesthetized with isoflurane (Viatris, VTRS) and subsequently perfused transcardially with 10 mL of 1 x phosphate-buffered saline (PBS) followed by 30 mL of 4 % paraformaldehyde (PFA) in 1 x PBS. The brains were extracted and post-fixed in 4 % PFA overnight at 4 °C, followed by a further 2-3 days in 30 % sucrose/PBS at the same temperature. The brains were then embedded in O.C.T. compound (Sakura Finetek Japan) and frozen at -80 °C. The brains were sectioned into 40 μm-thick coronal slices using a cryostat (Leica, CM1850 or Microm, HM505E). The expression of the virus and the locations of GRIN lens or fiber tips were verified using a fluorescence microscope (Keyence, BZ-X700 or BZ-X800). A confocal microscope (Oxford Instruments, Dragonfly) was used for the observation of terminal virus expression at a higher resolution. Images of axon terminals were captured using tiling and z-stack functions with a 20x objective. All animals used in the imaging experiments and experimental animals (injected with AAV5-hSyn-Chrimson-TdT or AAV5-hSyn-Jaws-KGC-GFP-ER2) used in the optogenetic experiments were excluded from the analyses if the GRIN lens or fiber placement was outside of the target area or virus expression could not be confirmed.

### Bayesian inference model

Here we summarized the key components of the model; full derivations, notation, and algorithm are in the Supplementary Methods.

#### Model formulation

We assumed that mice infer a latent contextual state *s*_*t*_ ∈ {aversive,safe} from observations *o*_*t*_ ∈ {shock,no shock} at each time step *t*. The latent context is the cause of observations: in aversive conditioning and extinction, the true cause is whether a stimulus is presented or omitted, and mice encode this as beliefs about an aversive or safe context. We formalized this inference as a POMDP with active inference, in which mice update contextual beliefs from observations (perceptual inference) and steer belief transitions via a policy *u*_*t*_ ∈ {to aversive,to safe} (active inference):

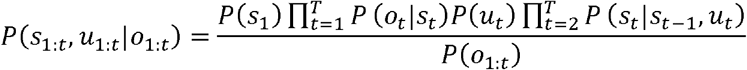

Where 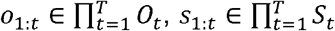, and 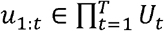 denote sequences of observations, contexts, and policies within a trial. Policies *u*_*t*_ formalize the mouse’s active steering of its own contextual beliefs.

We hypothesized that active inference operates during transitions from conditioning to extinction. Conventional latent cause models need strong prior assumptions to explain cautious belief updating. Active inference instead couples belief updating to policy selection, so that the pace of contextual inference emerges from goal-directed control rather than ad hoc biases. This accounts for non-canonical hippocampal activity patterns that are difficult to explain under existing theories.

To infer the true cause of observations, mice form approximate beliefs *Q*(·) bout contexts and policies:

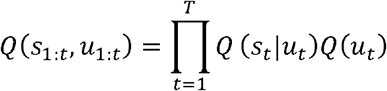

The objective is to match *Q*(·) to the generative model *P*(·) by minimizing the Kullback-Leibler divergence:

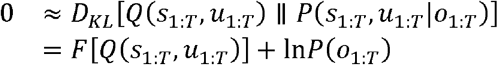

where

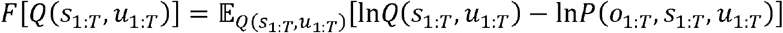

Minimizing this variational free energy *F*[·] improves the accuracy of the mouse’s predictions about observations.

#### Latent state inference and perception

Mice form contextual beliefs s_*t*_ by finding contexts *s*_1:*T*_ that minimize *F* at each time step. Decomposing *F* along the temporal direction and taking the gradient with respect to *s*_*t*_ under sophisticated inference gives:

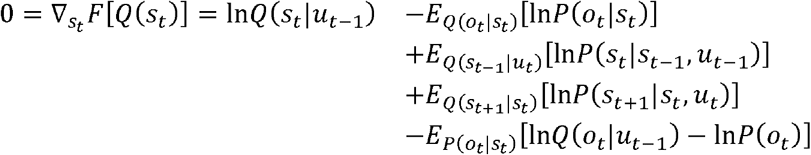

Each term in this equation can be expressed using sufficient statistics with parameters from the generative model *P*(·). The mouse’s belief models *o*_*t*_ = *p*(*O*_*t*_), s_*t*_ = *p*(*S*_*t*_), u_*t*_ = *p*(*U*_*t*_) are expressed as categorical distributions *p*(·) = Cat(·), parameterized by:

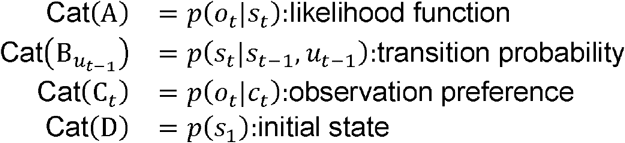

The transition matrix 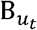 differs by selected policy *u*_*t*_, allowing mice to steer contextual transitions. Substituting these sufficient statistics into the gradient equation above and applying the sigmoid function *σ* (·)yields:

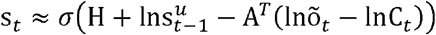

where 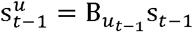 is the policy-dependent state prediction 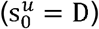, H = diag(A · lnA) is the entropy of the likelihood matrix, and õ_*t*_ is the target observation.

#### Active inference and freezing behavior

In active inference, mice prepare different transition probabilities 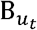 depending on the selected policy *u*_*t*_. Mice can therefore actively form contextual beliefs that better predict future observations, not just passively update from what they observe.

We expressed such optimal future contextual beliefs as:

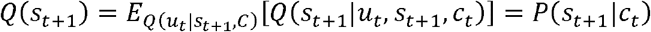

That is, a policy *u*_*t*_ infers a desirable context *P*(*S*_*t*+1_|*C*_*t*_) at the next time point. The mouse’s task is to form a policy *Q* (*u*_*t*_) that generates *Q*(*S*_*t*+1_) satisfying the above. This is achieved by selecting a policy that minimizes *expected free energy G (u*_*t*_):

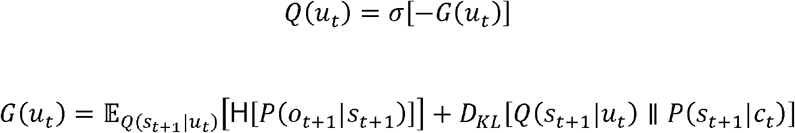

As with variational free energy, this can be written in terms of the generative model parameters:

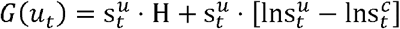

where 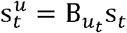 is the context after policy-driven transition and 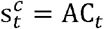 is the context that yields preferred observations.

We assumed mice exhibit freezing when they believe the current context shifts toward aversive state. At this time, the relationship between the mouse’s freezing response *fb*_*t*_ ∈ {freezing,non-freezing} and policy *u*_*t*_ is expressed as:

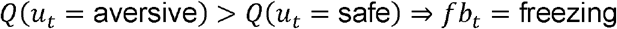

When such a relationship holds, freezing can be concisely expressed using a softmax function:

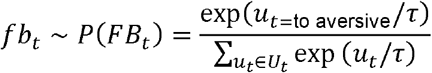

where *τ* is a temperature parameter (*τ* = 0: deterministic; , *τ* = ∞: random).

#### Modeling extinction learning

Extinction learning involves inferring the probability that electrical shock occurs after CS presentation. During the CS phase, mice observe sound rather than shock; we model the belief that sound predicts shock as C_*SS*_ = *P*(shock|sound) = Dir(c_*SS*_), a learnable component of the observation preference C_*t*_. Under variational inference, c_*SS*_ is updated on each trial (exact inference is also possible). To link extinction learning to the mouse’s epistemic process, we replaced the shock observation with contextual beliefs s_*t*_ during the shock phase:

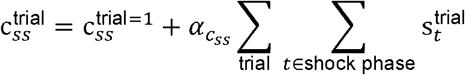

The rate of contextual inference during the post-CS phase is thereby coupled to the rate of extinction learning during the CS phase.

#### State prediction error and control error

We assumed that vCA1 neurons encode contextual beliefs s_*t*_. The sigmoid function *σ(·)* acts as an activation function (neural activation–firing rate relationship). The log-expectation lns_*t*_ is the neural activation of vCA1 neurons encoding context *s*_*t*_, which we denote *v*_*S,t*_.

The *state prediction error* ε_*S,t*_ is the temporal change in neural activation:

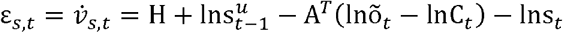

This corresponds to the negative gradient of variational free energy:

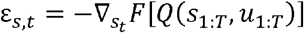

We assumed that context *s*_*t*_ is inferred via gradient descent on *F*. The update rule is:

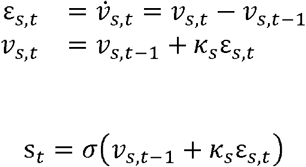

where *κ*_*S*_ is the learning rate. The integrated signal projected from vCA1 is *ε*_*t*_ *= ε*_*S*=aversive,t_ - *ε*_*S*=safe,*t*_; positive values promote updates toward the aversive context, negative values toward the safe context.

Beyond this perceptual inference error, we formalized projection signals from vCA1 as a post hoc evaluation of the selected policy. In the Bayesian network of the generative model (**Fig. 3a**), context s_*t*_ transmits information to policy u_*t*_; we assumed that vmPFC, NAc, and BA are associated with policy beliefs, i.e., active control of contextual transitions. We defined a *control error* as the policy-dependent deviation between current contextual beliefs and desirable contextual states, by factoring the negative expected free energy – *G*(*u*_*t*_):

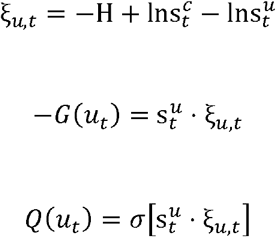

Policy inference is the product of the transitioned context 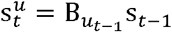 and the weighting factor ξ_*u, t*_.

Let 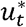 denote the policy selected at the previous time step. To find the optimal target context, we differentiated 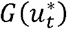 with respect to 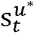 and set the gradient to zero:

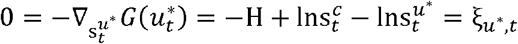

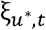 quantifies how far the selected policy displaced contextual beliefs from the desired context. State prediction error drives belief updating proportional to surprise (a perceptual inference error), whereas control error evaluates whether the selected policy moved the system closer to a desirable state (an active inference error). As with s_*t*_, policy *u*_*t*_ is updated via gradient descent. The integrated control error is:

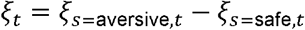

When negative, policy control toward the safe context is still needed; when positive, control toward the aversive context is needed.

#### Modeling optogenetic manipulation

Both *ξ*_*t*_ and *ξ*_*t*_ predict freezing (negative values correspond to high aversive belief), so correlational results alone cannot distinguish which signal vCA1 projections carry. To dissociate them, we embed state prediction error within control error. Using 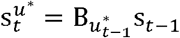:

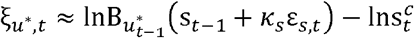

At the onset of the post-CS phase, s_*t* − 1_ is strongly aversive while 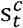 favors the safe context. Amplifying ε_*S,t*_ (making it more negative) accelerates the safe-context transition in the first term, reducing 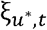. State prediction error and control error therefore have opposing effects on extinction under artificial manipulation: increasing the error signal impedes extinction if it is a state prediction error (larger error pushes updating away from current belief) but facilitates extinction if it is a control error (reducing the deviation from the desirable context). In simulations, optogenetic manipulation was modeled as:

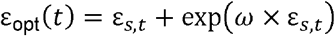

where *ω* is the manipulation strength.

### Statistical Analysis

Statistical analysis was performed with Prism 7 (GraphPad) and Matlab (Mathworks). Violin plots represent data distributions, and their lines represent the median and 25th and 75th percentiles. Box plots represent the median and 25th and 75th percentiles, and their whiskers represent the data range. Group differences were detected using Friedman test, Kruskal-Wallis test, or two-way repeated measures (RM) ANOVA where appropriate. Significant group effects in the Friedman test or Kruskal-Wallis test were followed by Dunn’s multiple comparisons test. Significant main effects or interactions in the ANOVA were followed by Holm-Sidak post hoc method. For paired comparison, we used Wilcoxon signed-rank test or paired t-test. All data analyzed by paired t-test were normally distributed according to Shapiro-Wilk test. All miniscope results were replicated across multiple cells of multiple animals. All photometry, behavioral, and model results were replicated across multiple animals. Throughout the study, P < 0.05 was considered statistically significant. No statistical methods were used to predetermine sample sizes, but our sample sizes are similar to those generally used in the field.

## Acknowledgement

We thank Drs. Joshua Johansen, Takaaki Ozawa, and Thomas McHugh for critical reading and comments on the draft; M. Maetani, K. Guzinsky, and R. Terakawa for technical assistance; and the members of Uematsu Lab for critical comments and discussion.

## Author contributions

A.S. and A.U. conceived the study. A.S., H.O., S.C.T. and A.U. developed the methodology. A.S., H.O., A.N.J., S.C.T. and A.U. carried out the investigation. K.E., A.S.T. and A.U. supervised the project. A.S., H.O., S.C.T. and A.U. wrote the original draft. A.S., H.O., K.E., and A.U. reviewed and edited the manuscript.

## Data availability statement

Custom code and full datasets will be made available upon request. The simulation model is available at: https://doi.org/10.5281/zenodo.18653800

**Supplementary Information is available for this paper**

## Notes

### Competing Interest Statement

The authors have declared no competing interest.

### Funding

- JST-FOREST JPMJFR2243 (A.U.) AMED-PRIME (A.U.) KAKENHI (20K21458, 21H05176, 22H02942) (A.U.) KAKENHI (16H06456, 16H02504) (K.E.) KAKENHI (JP21H05172) (S.C.T) KAKENHI (22K13838, 25K17180) (H.O.) KAKENHI (25KJ0934) (A.S.) JST SPRING (JPMJSP2108) (A.S.) Takeda Science Foundation (A.U., K.E.) AMED-CREST JP21gm310010 (K.E.) Torray Foundation (K.E.) Naito Foundation (K.E.) WPI-IRCN (A.U., K.E.)

