## Extended Data Figures for "A neural substrate for resistance to change in the ventral hippocampus"

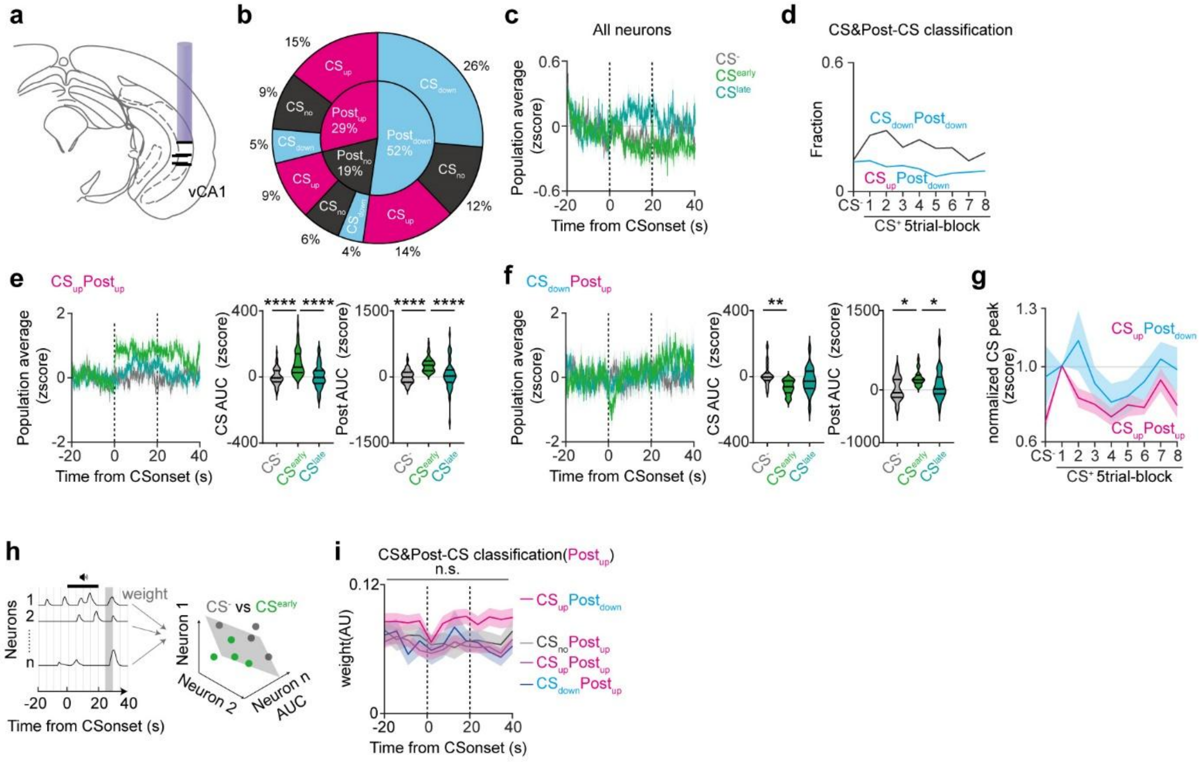

**Extended Data Fig. 1: Functional classification and response profiles of vCA1 neurons (Related to Figure 1).**

**(a)** Histological validation of GRIN lens implantation in vCA1 (n = 5). **(b)** Fractions of CS and Post-CS classes during $CS^{early}$. The majority of neurons showed inhibition in response to expected shock omission. **(c)** Population activity of all cells during $CS^{-}$, $CS^{early}$ and $CS^{late}$. Mean ± SEM. **(d)** CS&Post-CS classes across extinction. The fraction of $Post_{down}$ neurons gradually decreased across extinction session, which may reflect an extinction-associated switching. **(e)** Population activity of $CS_{up}Post_{up}$ cells during $CS^{-}$, $CS^{early}$ and $CS^{late}$(left). Mean ± SEM. Mean Z-score AUC of $CS_{up}Post_{up}$ cells during CS (middle) and post-CS (right) periods in extinction. CS and post-CS responses increased during $CS^{early}$ compared to $CS^{-}$. Lines, median, 25th and 75th percentiles. **(f)** Population activity of $CS_{down}Post_{up}$ cells during $CS^{-}$, $CS^{early}$ and $CS^{late}$(left). Mean ± SEM. Mean Z-score AUC of $CS_{down}Post_{up}$ cells during CS (middle) and post-CS (right) periods in extinction. CS responses decreased but post-CS responses increased during $CS^{early}$ compared to $CS^{-}$. Lines, median, 25th and 75th percentiles. **(g)** Mean peak of $CS_{up}Post_{up}$ and $CS_{up}Post_{down}$ cells during CS trial-blocks across extinction. Peaks were normalized to $CS^{early}$. Peaks of $CS_{up}Post_{down}$ were higher than $CS_{up}Post_{up}$ at late phase of extinction. **(h)** Schematic of population decoding using SVM. **(i)** Weights of $CS_{up}Post_{down}$, $CS_{up}Post_{up}$, and $CS_{down}Post_{up}$ cells. Weight of $CS_{up}Post_{down}$ cells was significantly higher than that of $CS_{up}Post_{up}$ or $CS_{down}Post_{up}$ at post-CS period. Mean ± SEM. n.s. not significant; *$P < 0.05$; For statistical details, see Table S1. *$P < 0.05$; **$P < 0.01$; ****$P < 0.0001$; For statistical details, see Supplementary Table 1.

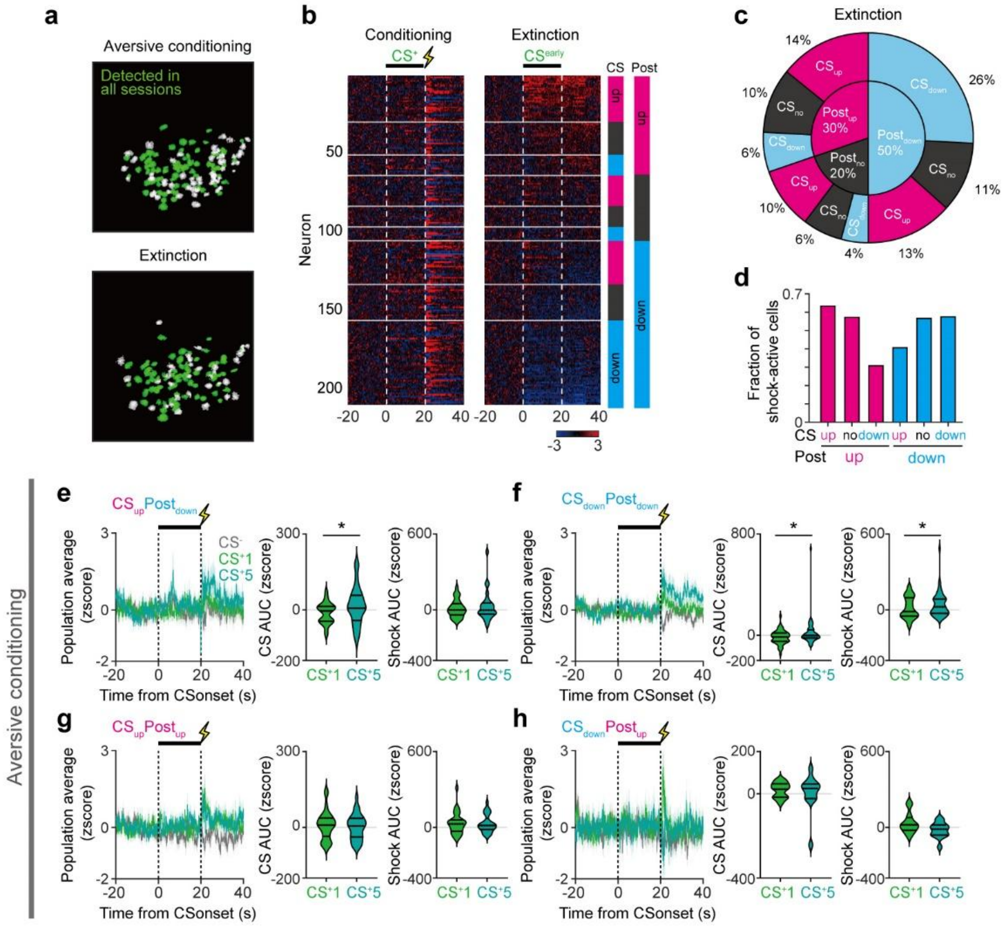

**Extended Data Fig. 2: Across-session imaging and response profiles of vCA1 neurons during aversive conditioning (related to Figure 1).**

**(a)** Registration of cells between aversive conditioning and extinction session. **(b)** Heat maps aligned to $CS^+$ onset during aversive conditioning (left) and $CS^{early}$ during extinciton (right) (n = 211 cells, 5 mice). Neurons are sorted based on $CS^{early}$ response categories during extinction, shown in magenta and blue squares. $CS_{up}Post_{up}$ (n = 30), $CS_{no}Post_{up}$ (n = 21), $CS_{down}Post_{up}$ (n = 13), $CS_{up}Post_{no}$ (n = 20), $CS_{no}Post_{no}$ (n = 13), $CS_{down}Post_{no}$ (n = 9), $CS_{up}Post_{down}$ (n = 28), $CS_{no}Post_{down}$ (n = 23), and $CS_{down}Post_{down}$ (n = 54) groups. **(c)** Fraction of functional cell types in longitudinal registration. **(d)** Fraction of shock-active cells within CS and post-CS groups. Statistical analysis revealed no significant differences across subgroups. **(e-h)** Population activity of $CS_{up}Post_{down}$ (e), $CS_{down}Post_{down}$ (f), $CS_{up}Post_{up}$ (g), and $CS_{down}Post_{up}$ (h) cells during $CS^-$, the first $CS^+$ trial ($CS^+1$) and the last trial ($CS^+5$) of aversive conditioning (left). Mean ± SEM. AUC of those cells during the CS (middle) and post-CS (left) period of $CS^+1$ and $CS^+5$. $CS_{up}Post_{down}$ cells developed CS responses during aversive conditioning (e). Also, $CS_{down}Post_{down}$ cells increased CS responses (f). Lines, median, 25th and 75th percentiles. $^*P < 0.05$; For statistical details, see Supplementary Table 1.

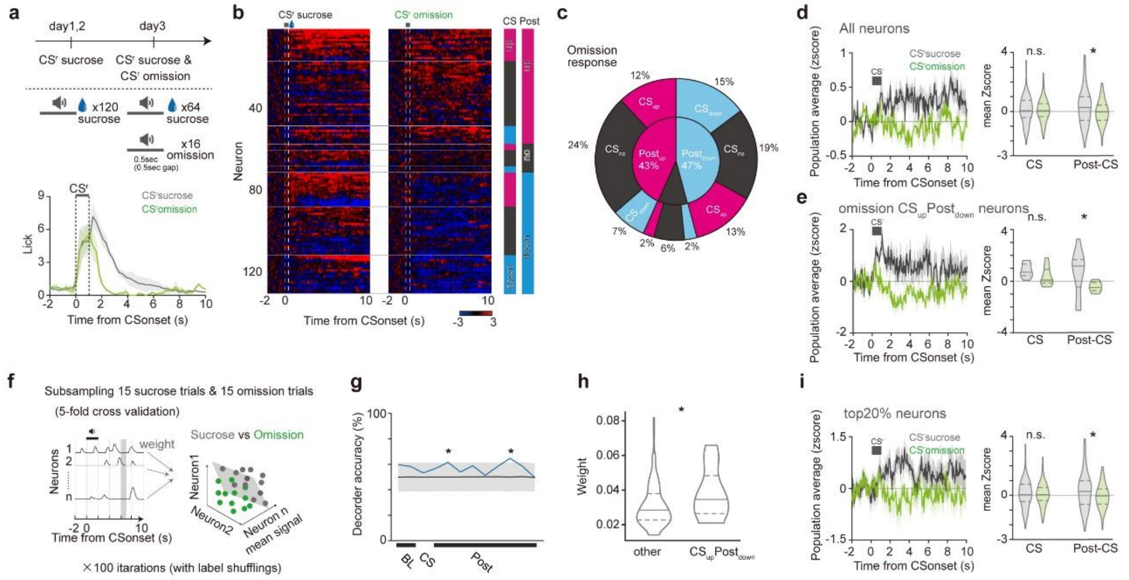


**Extended Data Fig. 3: vCA1 neurons exhibit reduced responses to the omission of expected appetitive outcomes. (related to Figure 1).**

**(a)** Reward conditioning and omission paradigm (top). Lick counts during sucrose and omission trials. Data represent group mean values (bottom). **(b)** Heat map during sucrose (left) or omission (right) trials (n = 131 cells, 3 mice). **(c)** Fractions of CS and Post-CS classes during omission. The majority of neurons showed inhibition in response to expected sucrose omission. **(d, e)** Population activity during sucrose and omission trials. The bar denotes the CS period (right). Mean signal of all cells at sucrose or omission trials (left). All neurons and $CS_{up}Post_{down}$ cells showed reduced Post-CS response during omission trials compared to sucrose trials. **(f)** Schematic of population decoding using SVM with subsampling comparable reward trials. **(g)** Time-resolved decoding accuracy of sucrose and omission trial-identity. Decoding accuracy exceeded that of label-shuffled data (gray line) at two post-CS points. **(h)** Weights of $CS_{up}Post_{down}$ and the other cells in linear decoder at the post-CS period, when decoding accuracy was significant. $CS_{up}Post_{down}$ cells showed higher classifier weights. **(i)** Population activity during sucrose and omission trials in top20% high-weight cells. The bar denotes the CS period (right). Mean signals during CS and Post-CS periods at sucrose or omission trials (left). These cells also exhibited reduced Post-CS response during omission trials compared to sucrose trials. n.s., not significant; *$P < 0.05$; For statistical details, see Supplementary Table 1.

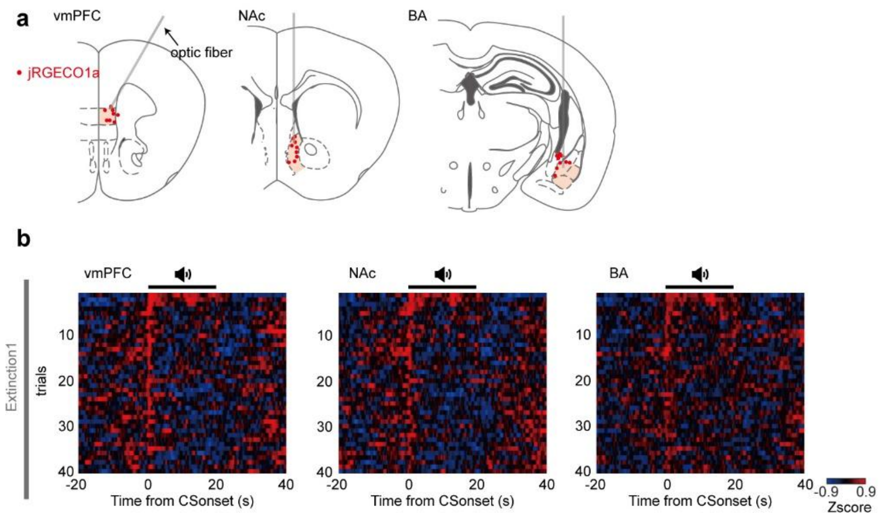


**Extended Data Fig 4: Trial-by-trial responses at individual axon terminals (related to Figure 2).**

**(a)** Histological validation of fiber implantation in vmPFC (left), NAc (middle), and BA (right). **(b)** Heat maps of each $CS^+$ trial of extinction session in vmPFC (left), NAc (middle), and BA (right) across all mice.

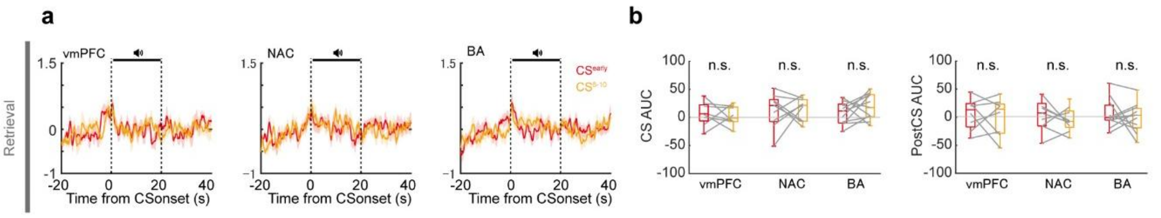


**Extended Data Fig. 5: Post-CS responses were diminished during extinction retrieval (related to Figure 2).**

**(a)** Mean terminal activities during CS$^{early}$ and 6-10 trials of CS$^{+}$ (CS$^{6-10}$) during retrieval. Small CS-evoked responses but no inhibition at post-CS period. Mean ± SEM. **(b)** Mean AUC during the CS and post-CS period of retrieval. No significant differences between CS$^{early}$ and CS$^{6-10}$. n.s. not significant; For statistical details, see Supplementary Table 1.

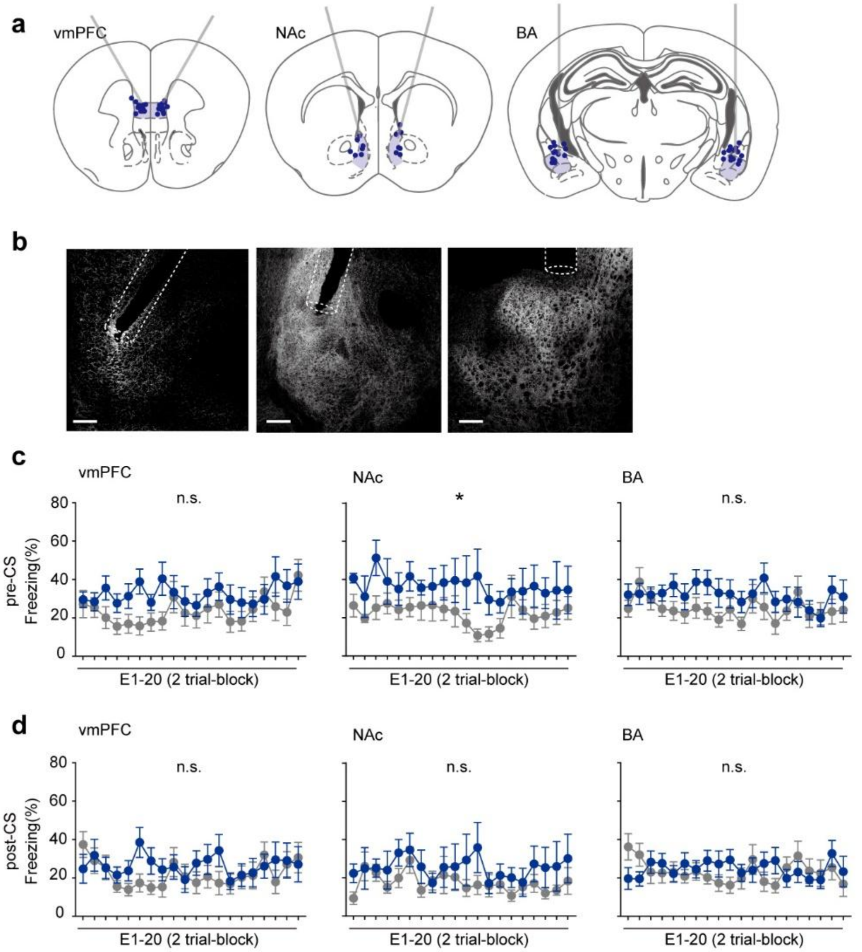


**Extended Data Fig. 6: Pre- and post-CS freezing levels in optogenetic inactivation (related to Figure 2).**

**(a)** Histological validation of fiber implantation in vmPFC (left, Jaws, n = 12), NAc (middle, Jaws, n = 7), and BA (right, Jaws, n = 14). **(b)** Representative Jaws expression and fiber implantation in vmPFC (left), NAc (middle), and BA (right). Scale bar, 200 μm. **(c, d)** Freezing behavior during the pre-CS (c) and post-CS (d) periods in inhibition (blue) and control (gray) groups. Optogenetic inhibition of NAc terminals increased pre-CS freezing. Mean ± SEM. n.s. not significant; **P* < 0.05; For statistical details, see Supplementary Table 1.

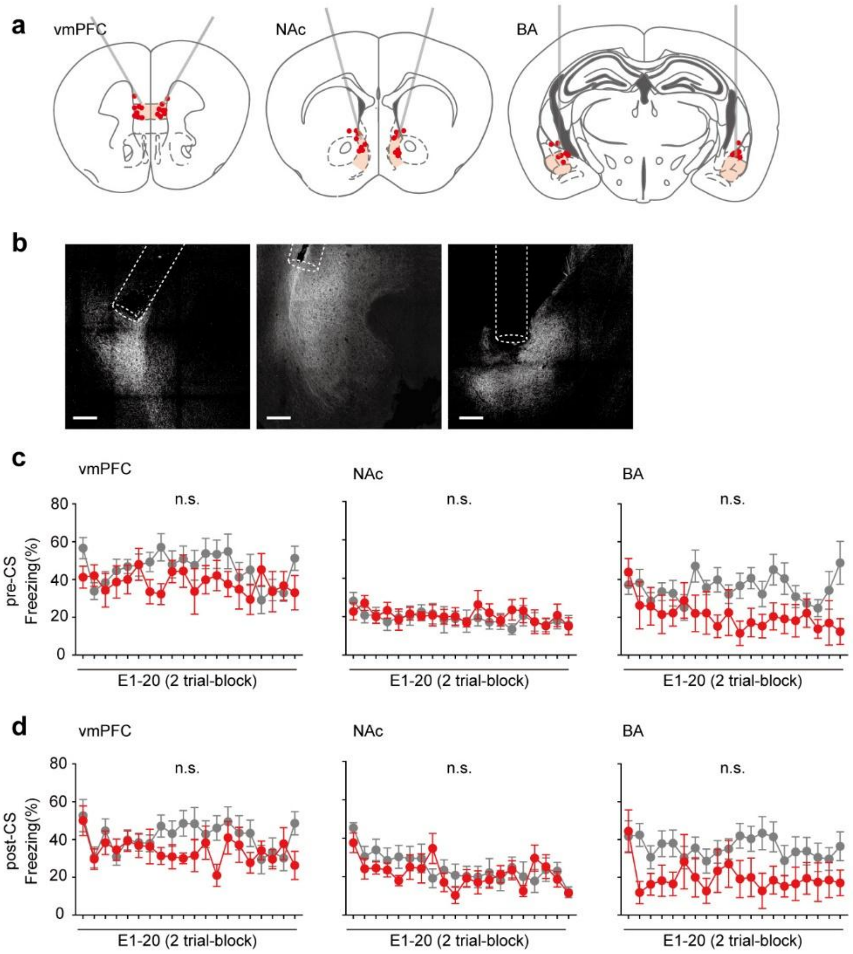


**Extended Data Fig. 7: Pre- and post-CS freezing levels in optogenetic activation (related to Figure 2).**

**(a)** Histological validation of fiber implantation in vmPFC (left, Chrimson, n = 8), NAc (middle, Chrimson, n = 8), and BA (right, Chrimson, n = 7) **(b)** Representative Chrimson expression and fiber implantation in vmPFC (left), NAc (middle), and BA (right). Scale bar, 200 µm. **(c, d)** Freezing levels during the pre-CS (c) and post-CS (d) periods in activation (red) and control (gray) groups. Optogenetic activation didn't affect pre- or post-CS freezing. Mean ± SEM. n.s. not significant; For statistical details, see Supplementary Table 1.

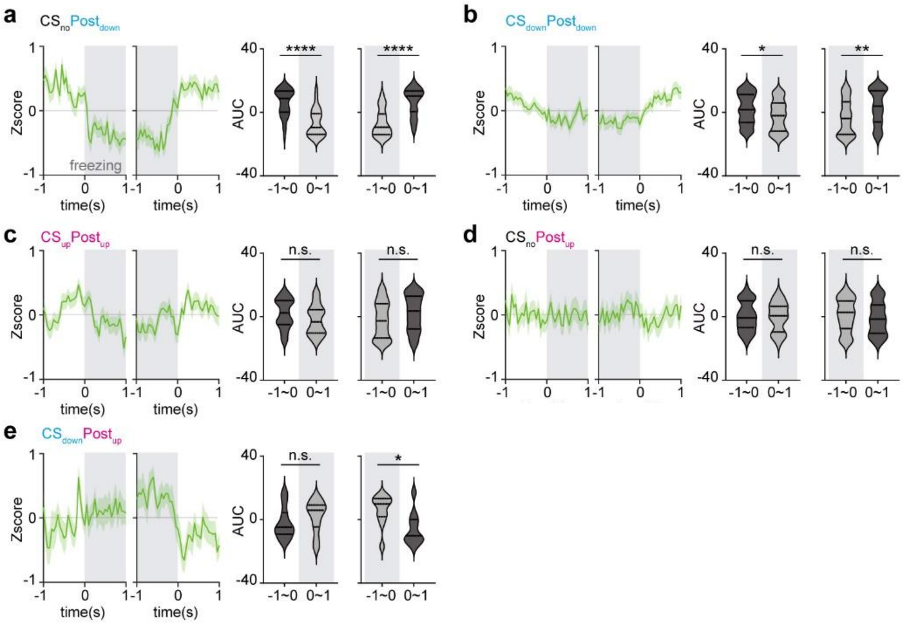


**Extended Data Fig. 8: Freezing-aligned activity across the functionally-defined cell types (related to Figure 4).**

**(a-e)** Population activity of CS and post-CS subclasses aligned to post-CS freezing onset and offset (left) in single cell experiments. Gray area indicates freezing period. $CS_{no}Post_{down}$ (a) and $CS_{down}Post_{down}$ (b) cells reduced responses after freezing onset and before freezing offset. $CS_{down}Post_{up}$ reduced responses after freezing offset (e). $CS_{up}Post_{up}$ (c) and $CS_{no}Post_{up}$ (d) responses were not affected by freezing. Lines in violin plots, median, 25th and 75th percentiles. Mean ± SEM. n.s. not significant; *$P$ < 0.05; **$P$ < 0.01; ****$P$ < 0.0001; For statistical details, see Supplementary Table 1.

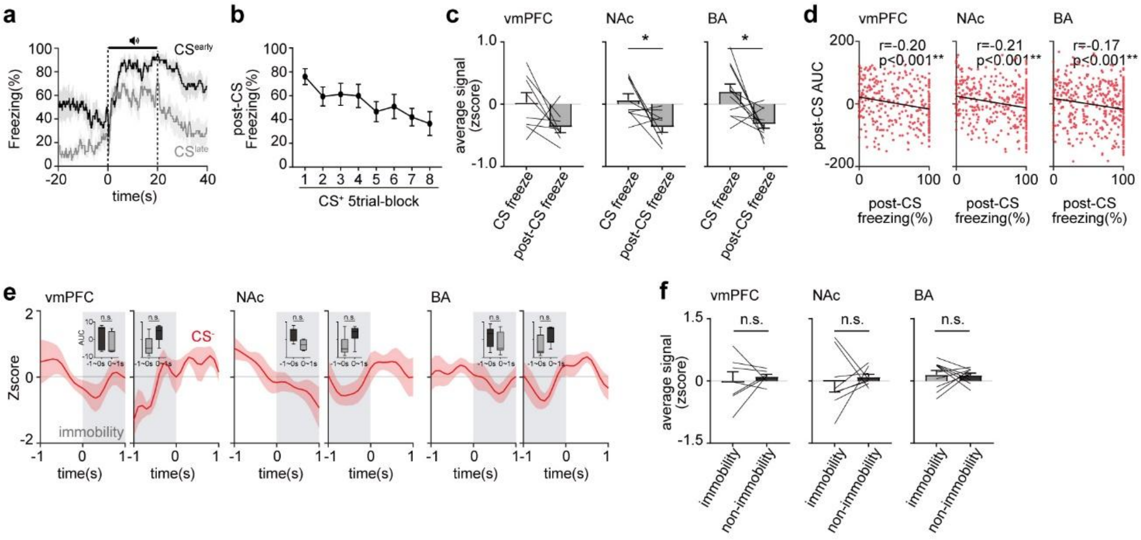


**Extended Data Fig. 9: Freezing- and immobility-aligned activity in multifiber photometry (related to Figure 4).**

**(a)** Time-resolved freezing levels during $CS^{early}$ and $CS^{late}$ of extinction1 across all mice. **(b)** Freezing levels during the post-CS period of extinction1 in multifiber photometry. Mean ± SEM. **(c)** Mean terminal activities during the CS or post-CS freezing period of $CS^{early}$ of extinction1 in vmPFC (left), NAc (middle), and BA(right). Mean ± SEM. **(d)** Correlation between freezing levels and AUCs during the post-CS period of extinction1 session in vmPFC (left), NAc (middle), and BA (right). All terminal activities were negatively correlated with freezing during post-CS period. **(e)** Mean Z-scored activity aligned to onset and offset of immobility (1sec) during the $CS^{-}$ period of aversive conditioning. Insets display mean AUC during 1sec before and after immobility onset and offset. No significant differences were observed. Mean ± SEM. **(f)** Mean activity during entire immobility or non-immobility period of $CS^{-}$ in aversive conditioning. No significant differences were observed in vmPFC, NAc, and BA. These results indicate that freezing-specific, but not immobility-specific, reduction in vCA1 terminal activity. Mean ± SEM. n.s. not significant; **P* < 0.05; ***P* < 0.01; For statistical details, see Supplementary Table 1.

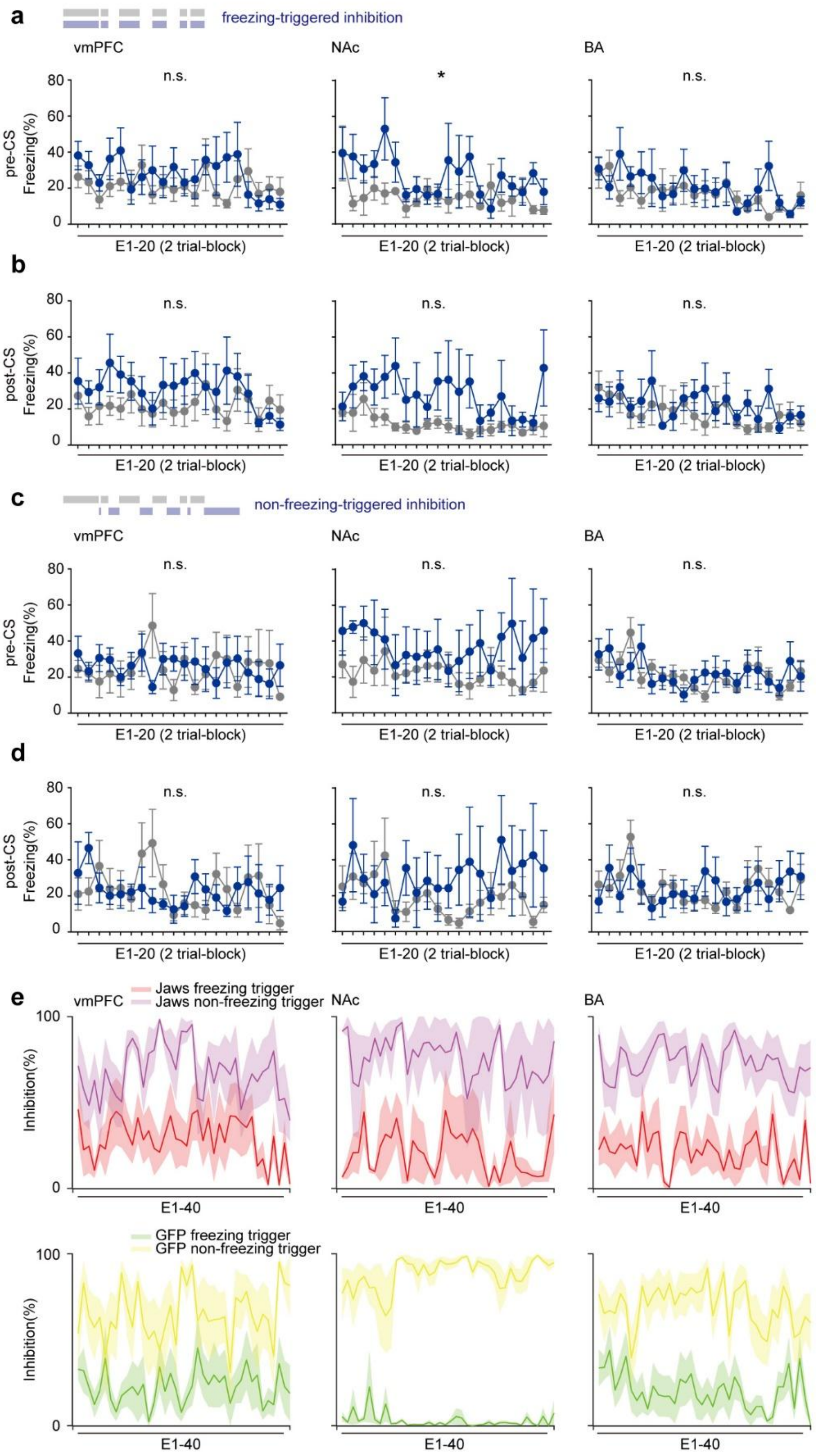


**Extended Data Fig. 10: Pre- and post-CS freezing levels in closed-loop optogenetic inactivation (related to Figure 4).**

**(a-d)** Freezing levels during the pre-CS (a, c) and post-CS (b, d) periods of extinction in closed-loop optogenetic inhibition. Blue and gray lines represent Jaws and control, respectively. Freezing-triggered (a, b) or non-freezing-triggered (c, d) groups. Freezing-triggered inhibition at NAc terminals during post-CS period increased freezing during pre-CS, consistent with entire post-CS inhibition (Extended Data Fig. 6).
**(e)** Inhibition duration of freezing-triggered and non-freezing-triggered conditions in Jaws (top) and control (bottom) groups. The duration of post-CS laser delivery was significantly shorter in freezing-triggered group than in non-freezing group, indicating that the observed effect on freezing in freezing-triggered group was not due to prolonged inhibition. Mean ± SEM. n.s. not significant; *$P < 0.05$; For statistical details, see Supplementary Table 1.
